# Genetic regulation of homeostatic immune architecture in the lungs of Collaborative Cross mice

**DOI:** 10.1101/2021.04.09.439180

**Authors:** Brea K. Hampton, Kara L. Jensen, Alan C. Whitmore, Colton L. Linnertz, Paul Maurizio, Darla R. Miller, Clayton R. Morrison, Kelsey E. Noll, Kenneth S. Plante, Ginger D. Shaw, Ande West, Ralph S. Baric, Fernando Pardo-Manuel de Villena, Mark T. Heise, Martin T. Ferris

**Affiliations:** Curriculum in Genetics and Molecular Biology, University of North Carolina at Chapel Hill, Chapel Hill, NC; Department of Genetics, University of North Carolina at Chapel Hill, Chapel Hill, NC; Department of Epidemiology, University of North Carolina at Chapel Hill, Chapel Hill, NC; Curriculum in Bioinformatics and Computational Biology, University of North Carolina at Chapel Hill, Chapel Hill, NC; Department of Microbiology and Immunology, University of North Carolina at Chapel Hill, Chapel Hill, NC; Lineberger Comprehensive Cancer Center, University of North Carolina, Chapel Hill, NC

**Author notes:** Co-senior/co-corresponding author **Mailing Address:** Department of Genetics, The University of North Carolina at Chapel Hill, 160 Dental Circle, Chapel Hill, NC, **Email:**.

**Keywords:** host genetics, homeostasis, lung immunity, quantitative trait loci, multiparent population

## Abstract

Variation in immune homeostasis, immune system stability, in organ systems such as the lungs is likely to shape the host response to infection at these exposed tissues. We evaluated immune homeostasis in immune cell populations in the lungs of the Collaborative Cross (CC) mouse genetic reference population. We found vast heritable variation in leukocyte populations with the frequency of many of these cell types showing distinct patterns relative to classic inbred strains C57BL/6J and BALB/cJ. We identified 28 quantitative trait loci (QTL) associated with variation in baseline lung immune cell populations, including several loci that broadly regulate the abundance of immune populations from distinct developmental lineages, and found that many of these loci have predictive value for influenza disease outcomes, demonstrating that genetic determinants of homeostatic immunity in the lungs regulate susceptibility to virus-induced disease. All told, we highlight the need to assess diverse mouse strains in understanding immune homeostasis and resulting immune responses.

## Introduction

Immune homeostasis is the stable state the immune system maintains in the absence of insult. While the majority of studies on host immunity have focused on the response to specific stimuli (e.g., pathogens, vaccines, allergens, or adjuvants), a growing body of evidence suggests that an individual’s baseline immune status affects subsequent innate or antigen specific immune responses (Gnjatic et al., 2017; Graham et al., 2019; HIPC-CHI Signatures Project TeamHIPC-I Consortium, 2017; Tsang et al., 2014). Immune homeostatic parameters are gaining increasing recognition as predictors of clinical outcomes to immunotherapy and vaccination (Gnjatic et al., 2017; HIPC-CHI Signatures Project TeamHIPC-I Consortium, 2017), and several studies have shown that dysregulation of immune homeostasis can contribute to the development of cancer, autoimmunity, allergies, as well as the progression of immune-related pathology in response to infection (Crimeen-Irwin et al., 2005). There is growing evidence that immune homeostasis is under genetic control (Graham et al., 2017; Krištić et al., 2018; Phillippi et al., 2013; Collin et al., 2019), however, the specific genetic factors that affect baseline immune function and cellular composition are poorly understood. In addition to aiding our understanding of the steady-state immune system, efforts to identify those specific genetic variants regulating homeostasis are likely to improve our understanding of how homeostasis affects later immune challenges or insults.

To date, much of the analysis of immune homeostasis has focused on the systemic immune system, such as total serum antibody or circulating immune cells in the blood or spleen (Cassidy et al., 1974; Graham et al., 2017; Grundbacher, 1974; 2018b). Less studied, but as important, is immune homeostasis in organ systems where the immune system routinely interacts with the environment. In particular, the lungs represent a major organ system that is constantly exposed to the external environment and as such are a common site of pathogen exposure. The lungs are protected by both immunological (e.g. resident immune cells and stromal cells) and non-immunological mechanisms (e.g. mucosal layer). The immune landscape in the lung has been shown to be influenced by age (Lloyd and Marsland, 2017; Marsland and Gollwitzer, 2014) and microbial exposure (Lloyd and Marsland, 2017; Jackson et al., 2008), with significant remodeling over time. Adding to these general processes are several studies having shown that microbial and environmental antigen exposure through the lungs can modulate subsequent infection (Blevins et al., 2014; Jackson et al., 2008), with many of these common environmental exposures linked to lung dysfunction later in life (Arrieta et al., 2015). Therefore, being able to expand our knowledge of lung-specific immune homeostasis to include genetic regulation, including specific genetic differences affecting homeostasis, will expand our understanding of the development of susceptibility to other pulmonary disease states.

While many studies have shown that humans exhibit significant inter-individual variation in baseline immune phenotypes (Cassidy et al., 1974; Grundbacher, 1974; HIPC-CHI Signatures Project TeamHIPC-I Consortium, 2017; Tsang et al., 2014), it has been particularly difficult to identify the genes that contribute to this variation due to a variety of confounding factors. As mentioned above, environmental factors, such as prior microbial and environmental exposures as well as aging differences create a highly dynamic immune environment in the lungs. Further, it is also physiologically challenging to perform invasive or mechanistic studies of homeostatic immunity in relevant tissues in humans. Rodent models represent an attractive system with which to model and investigate those factors driving the development of various immune homeostatic states. Specifically, the availability of reproducible inbred mouse strains and gene specific knockouts has allowed investigators to study how the presence of specific genes modulate both baseline and induced immunity across cellular compartments. These approaches have identified roles for genes like *IgHm* and *Rag1/Rag2* in immunity and immune development (Falk et al., 1996; Kitamura et al., 1991; Lansford et al., 1998), and *Foxp3* in immune regulation (Khattri et al., 2001; Kasprowicz et al., 2003). Gene specific knockout mice have been critical to our understanding of the immune system; however, these models represent extreme genetic perturbations, rather than the more subtle effects on gene expression or function more commonly associated with naturally occurring genetic variation in humans. Further, even in these model systems, most studies have analyzed the effects of genetic variation on baseline immune function in the systemic immune system (Orrù et al., 2013) (e.g. circulating immune cells or antibody) as opposed to compartment-specific homeostasis. As some immune cells populations are organ specific (e.g. alveolar macrophages, and the central nervous system-specific microglia), and immune cell composition and basal activation status is likely to vary from organ to organ, it is critical to understand compartment-specific genetic effects. Therefore, we sought to assess how genetic variation affects baseline immunity in the lung, a site of primary immune exposure, and further determine if this variation affects subsequent responses to pathogen exposure.

The Collaborative Cross (CC) genetic reference panel is an octo-parental set of recombinant inbred (RI) mouse strains derived from five classical laboratory strains (C57BL/6J (BL/6J), A/J, 129S1/SvImJ (129S1), NOD/ShiLtJ (NOD), and NZO/HlLtJ (NZO)) and three wild-derived strains (PWK/PhJ (PWK), CAST/EiJ (CAST), and WSB/EiJ (WSB)) (Collaborative Cross Consortium, 2012). These eight founder strains capture >90% of common genetic variation present in laboratory mouse strains and represent the three major subspecies of *Mus musculus* (Threadgill et al., 2011; Welsh et al., 2012; Roberts et al., 2007). We and others have shown that there is extensive variation in splenic T cell populations (Graham et al., 2017), antibody glycosylation patterns (Krištić et al., 2018), as well as variation in the general immune landscape of the spleen (Collin et al., 2019), across the CC population. We therefore sought to extend this work in the CC to investigate how genetic variation impacts immune homeostasis in the environmentally exposed lungs. Here, we took advantage of a panel of 95 genetically unique but reproducible CC-F1 hybrids (Noll et al., 2020), to assess the breadth of variation in immune cell populations in the lungs and attempt to identify loci regulating these phenotypes at the steady state to maintain immune homeostasis. We characterized a set of 54 cellular phenotypes spanning the innate and adaptive components of the immune system present in the unperturbed lung. We found that the immune cell populations we catalogued in the lung varied extensively (i.e. orders of magnitude in difference) between F1 hybrids; that genetic regulation of these immune cell populations was ubiquitous and strong; and further, we identified several genetic loci contributing to differences in their abundances. Several of these loci showed more extensive effects on the broader immune composition of the lung and also on disease in the context of Influenza infection. These results suggest that genetic regulation of immune homeostasis is a significant and critical component of the immune homeostatic environment, and that genetically distinct mouse models can aid in better understanding how genetic control of lung immune architecture can drive responses to immune insults and disease responses.

## Results

### Homeostatic immune landscape in the lungs of genetically diverse CC-F1 hybrids

To understand the role of genetic variation on lung leukocyte frequency in the absence of immune challenge, we evaluated lung leukocyte composition in 658 mice representing 95 distinct CC-F1 hybrids (3-9 female mice/line, 8-12 weeks old). Mice were sham challenged with PBS, as one arm of a larger study (Graham et al., 2017; Noll et al., 2020). Lungs were harvested at four days post PBS instillation and processed for flow cytometric analysis. The lungs have several different immune cell populations, including resident, inflammatory, and circulating leukocytes, and we assessed cell types from all of these categories (Table 1). We observed high levels of variation across the CC-F1 panel in all cell populations catalogued, as well as immunologically relevant meta-phenotypes, such as the ratio of specific cell populations (e.g. CD4:CD8 T cells). As previous studies of systemic immune populations have shown, T-cell populations in the lung are highly variable. CD4^+^ T cells ranged from 1.0 – 98.4% of CD3^+^ T cells in the lungs (Figure 1A). CD8^+^ T cells showed a similar pattern where 0.12 – 58.5% of CD3^+^ T cells represented this population (Figure 1B). Additionally, myeloid cell populations traditionally found in the lungs and tissue-resident immune cells were also variable across CC-F1s. Ly6C^-^‘patrolling’ monocytes varied from 0.08 – 26.4% of all LCA^+^ cells, plasmacytoid DCs ranged from 0 – 9.3% of LCA+ cells, and alveolar macrophages ranged from 0.1 – 35.8% of LCA+ cells in the lungs (Figure 1C, 1D, 1E). Lastly, populations of immune cells that are thought to primarily invade the lung under inflammatory conditions were highly variable across our CC-F1 population. For example, we found that eosinophils ranged from 0 – 41.9% of all LCA^+^ cells, neutrophils ranged from 0 – 56.6% of LCA^+^ cells, and Ly6C^hi^ ‘inflammatory’ monocytes/macrophages ranged from 0 – 26.3% of LCA+ cells in the lungs (Figure 1F, 1G, 1H). Thus, regardless of the origin or role of various immune cell populations there was vast variation in the levels of all cell populations measured across our CC-F1 panel.

**Table 1:**
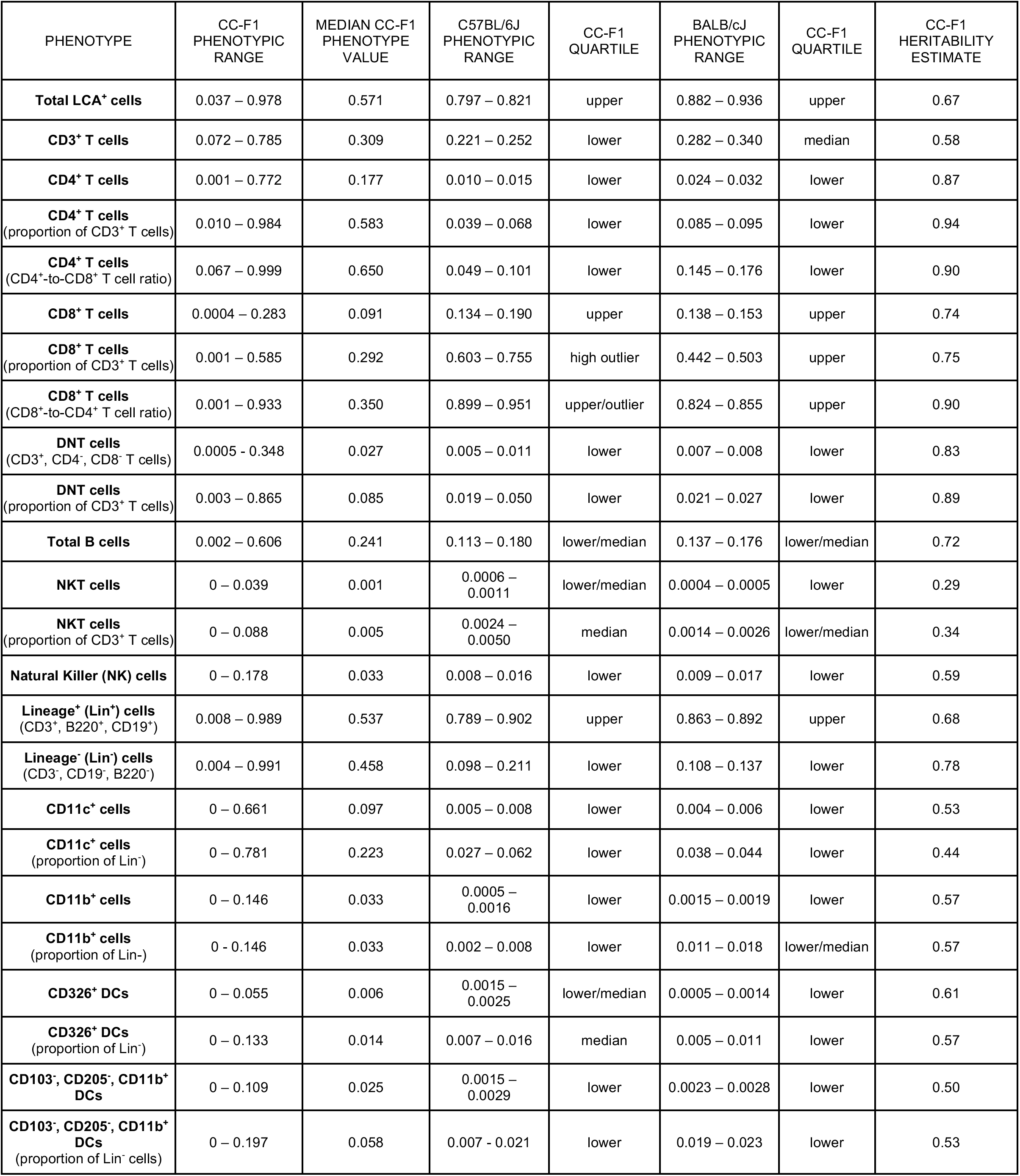
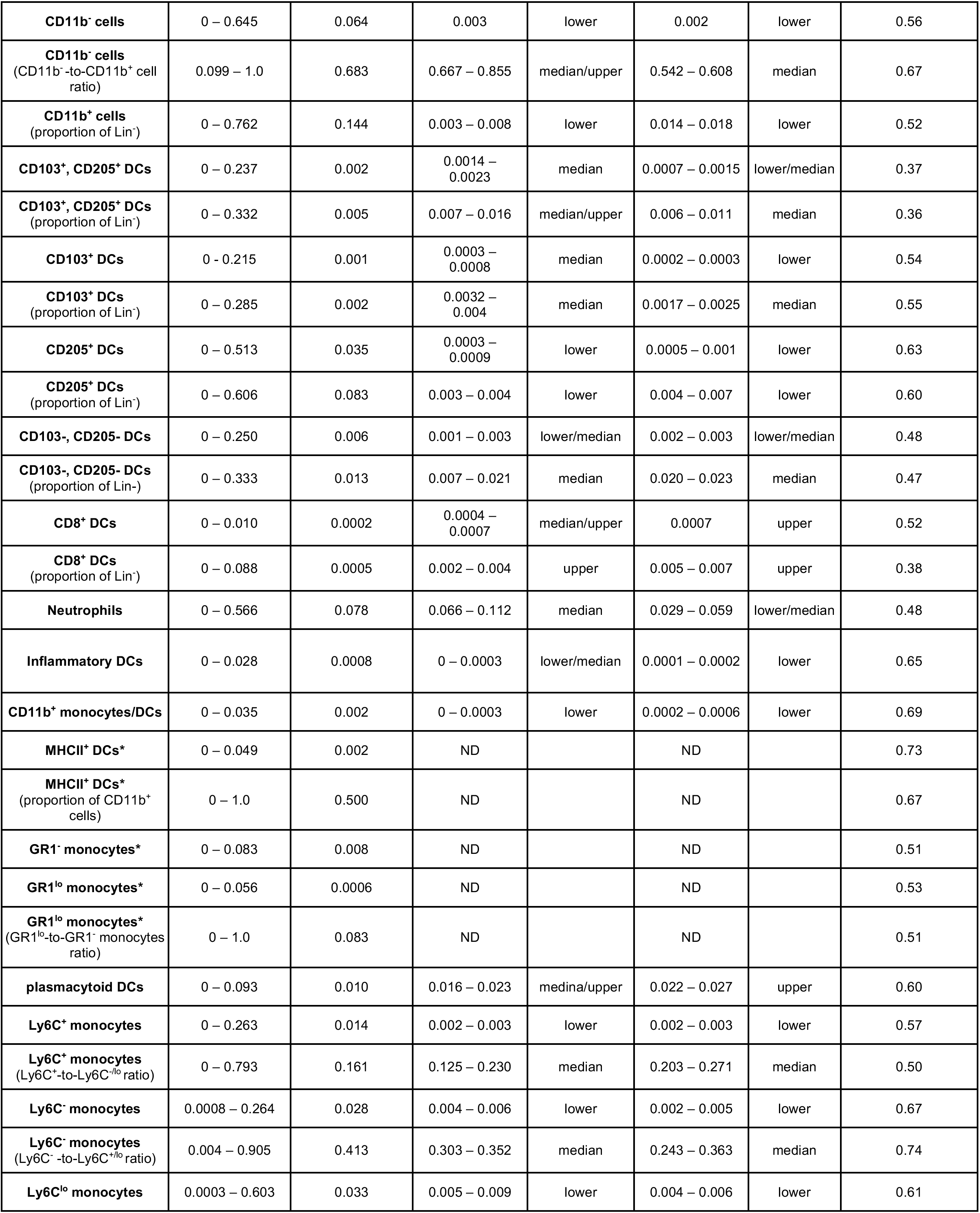
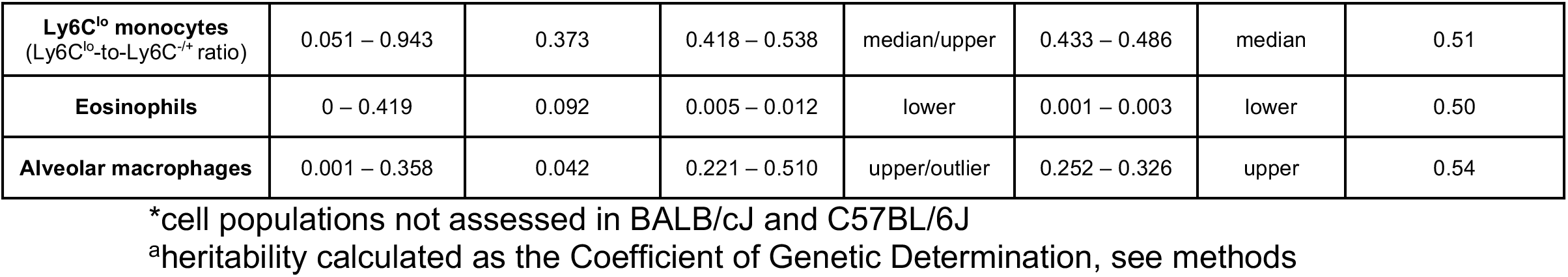
Cell populations measured, phenotypic ranges (proportions) and heritability estimates for CC-F1 hybrids and comparisons to classic BL/6J and BALB/cJ strain values.

**Figure 1:**
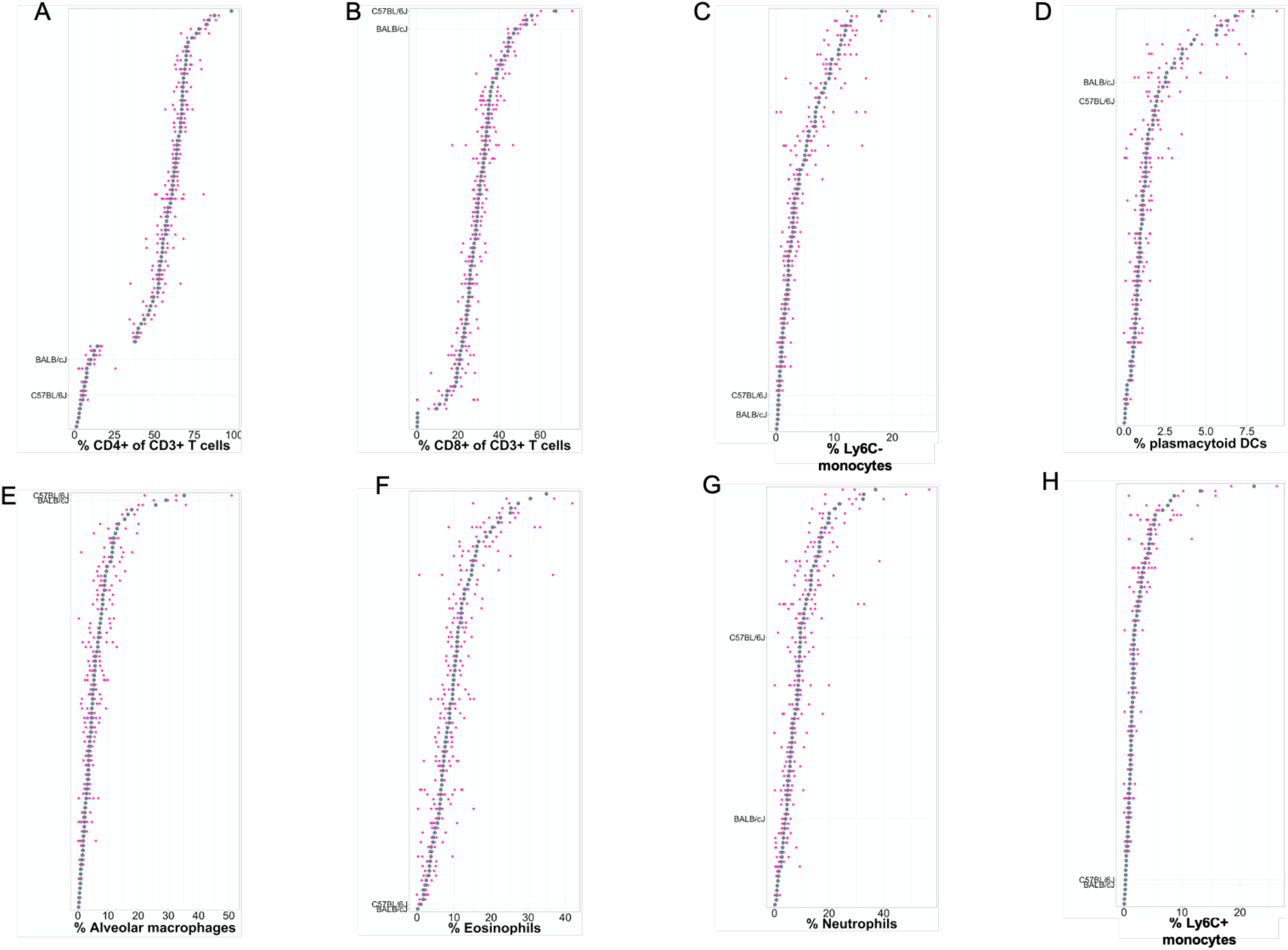
Homeostatic immune landscape in the lungs of genetically diverse CC-F1 animals. The mean for C57BL/6J, BALB/cJ, and each CC-F1 cross is shown in grey and the measurement for each individual animal within that cross are plotted in magenta, showing the variation around the mean. Phenotype distributions, where strains are independently ordered on the y-axis by the percentage of **A**) CD4+ T cells (% of CD3+ T cells), **B**) CD8+ T cells (% of CD3+ T cells), **C**) Ly6C^-^ monocytes, **D**) plasmacytoid DCs, **E**) alveolar macrophages, **F**) eosinophils, **G**) neutrophils, **H**) Ly6C^hi^ monocytes. On the y-axis, each CC-F1 or inbred strain is independently ordered within each phenotype. The x-axis varies between each phenotype and denotes the range of each phenotype.

Given this observed phenotypic variation, we estimated the broad sense heritability (proportion of trait variance attributable to genetic variation) for each cell population. We found that broad sense heritability estimates for the measured phenotypes in our CC-F1 panel ranged from 0.29 – 0.94 (median heritability = 0.57, Table 1), indicating variable but significant levels of genetic control of these traits. We next assessed the relationship between cell populations by assessing pairwise correlations between all cell populations (Figure 2A). While the majority of pairs show no to weak correlations (2400/2916, mean ρ = 0.049, range = 0.3163 > ρ > −0.2181), a subset of populations showed strong correlations (134/2916 comparisons, ρ > 0.58 or < −0.48). As expected, we find negative correlations such as the well described one between CD4^+^ T cells and CD8^+^ T cells as a proportion of total CD3^+^ T cells in the lungs (Figure 2B, ρ = −0.2759), and also a strong negative correlation between CD4^+^ T cells and DN T cells (Figure 2C, ρ = −0.9143). However, we also found several correlations not necessarily expected based on known immunological development. For example, we found a positive correlation between NK cells and Ly6C^lo^ monocytes/macrophages (ρ= 0.5636, Figure 2E). Interestingly, we find what appears to be a mutually exclusive relationship between DN T cells (CD4^-^, CD8^-^) and MHC II^+^ DCs (Figure 2D). All told, these relationships suggest that genetic coregulation between some of these cell populations is at play.

**Figure 2:**
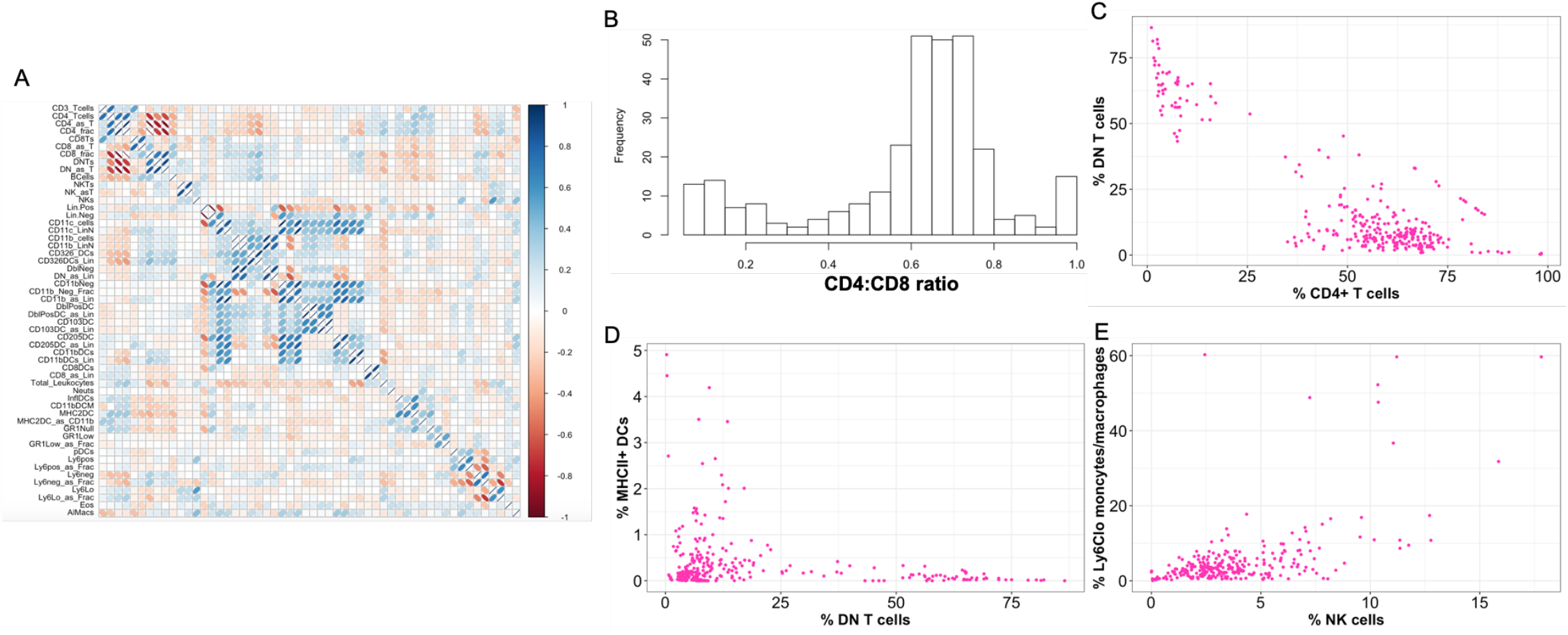
Lung immune cell correlation structure in CC-F1 hybrids. A) Per-animal pairwise correlations (x-axis is mirrored y-axis) were calculated between each cell population. The strength and direction of correlations are illustrated with both color (blue = positive, red = negative) and shape (more defined shapes are stronger correlations). B) Histogram of CD4:CD8 T cell ratios using individual animal measurements. C) Negative correlations between CD4^+^ T cells and DN T cells (correlation coefficient = −0.9143) in the lungs of CC-F1 mice. D) Mutually exclusive relationship between DN T cells and MHCII^+^ DCs in the lungs of CC-F1 mice. E) Positive correlation between NK cells and Ly6C^lo^ monocytes/macrophages (correlation coefficient = 0.5636) in the lungs of CC-F1 mice. Each pink dot (C-E) represents an individual animal.

BL/6J and BALB/cJ mouse inbred strains are frequently used in immunology research. We compared variation in immune cell populations across CC-F1 hybrids with that observed in these two laboratory strains of mice by assessing the same immune cell populations in the lungs of BL/6J and BALB/cJ animals (Table 1). While the standard inbred strains were in the middle quartile of the CC-F1 distribution for 12 of our 54 cellular phenotypes measured, these common laboratory strains were more frequently (30 phenotypes) in an outlier quartile or more extreme (Table 1, Figure 1). For example, we found that neutrophil proportions in BL/6J and BALB/cJ lungs were in the median quartiles and fell within the distribution observed in CC-F1 animals (Figure 1D). However, in both common lab strains we found extremely low eosinophil proportions in the lungs, as compared to the CC-F1 population (Figure 1C). We also observed low frequencies of CD4+ T cells, and high frequencies of CD8+ T cells, where BL/6J animals exceeded the ranges observed in CC-F1 mice and BALB/cJ mice were in the upper quartile of the distribution (Figure 1A, 1B). Together, our data suggest that the BL/6J and BALB/cJ strains are frequent outliers with respect to their baseline lung immune profiles, and as such they do not represent the breadth of the homeostatic lung environment seen across laboratory mouse strains.

Given the breadth of phenotypic variation in the context of our minimally perturbed (PBS installation 4 days before assessment) cohort, we sought to determine whether genotype-specific differences in cell populations could be due to PBS instillation, and therefore not reflective of true homeostatic variation. We selected six CC strains, as well as the standard BL/6J and BALB/cJ strains to assess the relative contributions of genetics versus innocuous perturbation on immune cell populations. We measured 19 lung cellular phenotypes, with or without PBS instillation; representing a subset of all cellular phenotypes catalogued in our larger screen. For 15/19 populations (Sup. Table 1), we found no evidence that PBS altered population abundances. Four cellular phenotypes did show evidence of PBS related perturbation in the lungs, however these perturbations occurred across all strains (Sup. Table 1). Thus, while some cell populations may exhibit PBS perturbations, this effect impacted strains equivalently, and therefore, the data from our larger CC-F1 panel represents true homeostatic effects and genetic differences.

### Identification of genetic loci regulating immune homeostatic landscape in the lungs of CC-F1 hybrids

Strain-specific (i.e. genetic) differences accounted for between 29-94% of the total phenotypic variation observed in the cell populations measured in our study (Table 1). We sought to identify specific genomic loci where variants are associated with differences in the abundance of various immune cell populations in the lungs via quantitative trait locus (QTL) mapping. We identified 28 QTL (four genome-wide significant (p < 0.05), and 24 suggestive QTL (6 at p < 0.1; 19 at p < 0.2)) associated with variation in 24 cellular phenotypes and meta-phenotypes (summarized in Figure 3A, details in Supplemental Table 2). These cellular populations span the tissue-resident, inflammatory and lymphocytic compartments. Altogether, the 28 loci identified here spanned the range of phenotypes measured. For the 28 loci identified here, the effect sizes (percentage of phenotypic variance explained by a QTL) range from 0.32 – 32.5%, with a concurrent explanation of heritability (0.63-79.7%; both summarized in supplemental table 2). We further noticed two major trends across our mapped loci: 1) most loci (25/28) impacted only one phenotype (a range of 1-4 traits/locus, median = 1 trait); and 2) several traits (14/24) only had a single significant locus detected.

**Figure 3:**
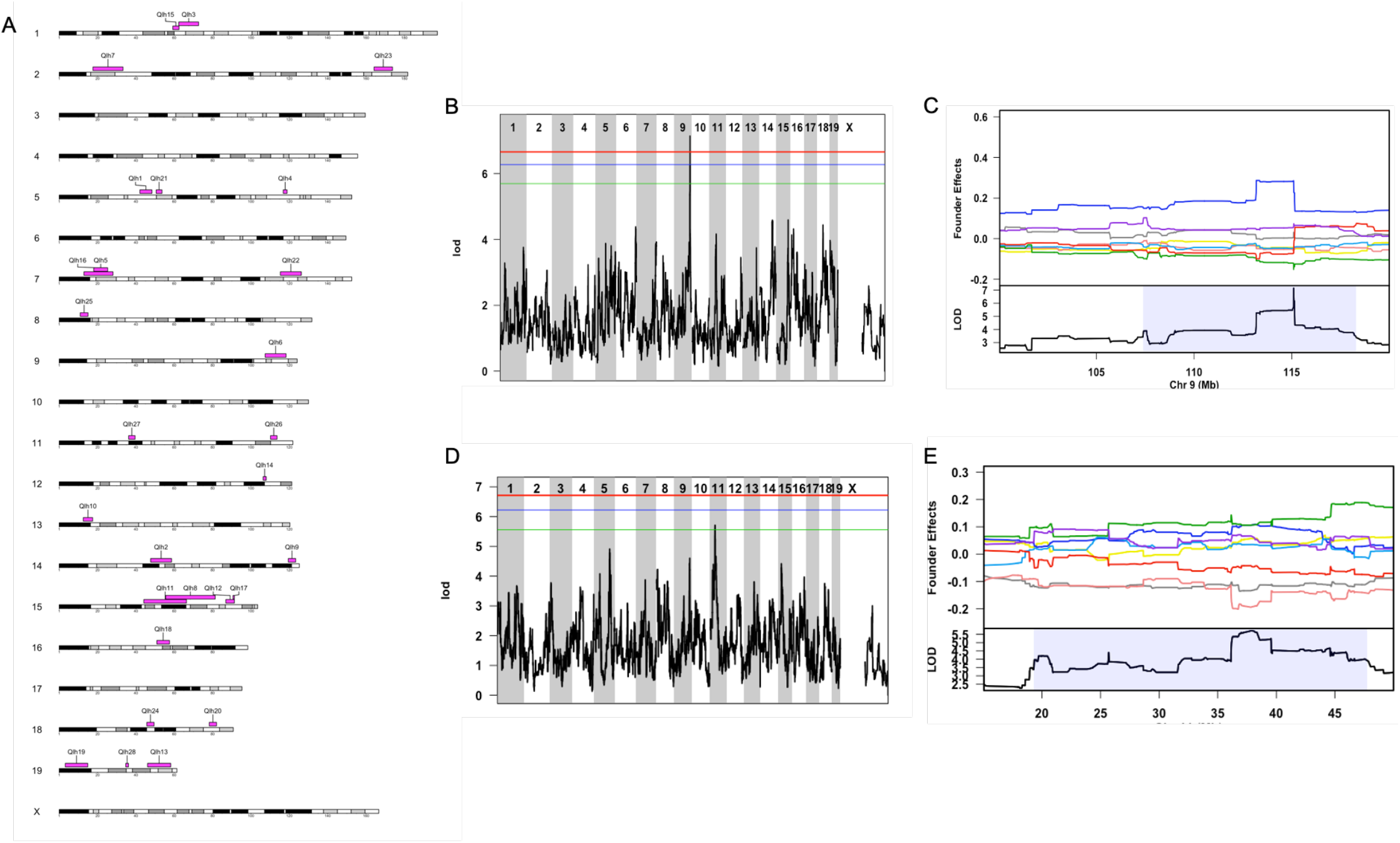
Immune landscape in the lungs of CC-F1 hybrid mice at the steady state is driven by multiple genetic loci. A) QTL contributing to lung immune cell composition are shown across the 20 mouse chromosomes (19 autosomes and X). Each QTL identified at a genome-wide p < 0.2 is shown at its position (pink blocks represent a 95% credible interval). Further details of the cell populations affected by these loci, and their allele effects are shown in Supp table 2). Representative *Qlh* QTL and allele-effect plots show distinct alleles contribute to immune homeostasis. LOD plots (panels B (alveolar macrophages) and D (neutrophils)) show QTL significance (y-axis) across the genome (x-axis) with significance thresholds (genome-wide p-value = 0.05 (red), 0.1 (blue), 0.2 (green)). Identified QTL had their associated allele effects (A/J = yellow, BL/6J allele = grey, 129S1 allele = pink; NOD allele = dark blue; NZO allele = light blue; CAST = green; PWK = red; WSB = purple) determined for the associated peaks C) alveolar macrophages and E) neutrophils. Allele effect plots show the mean deviation from population-wide mean as on the upper Y-axis for each allele segregating in the CC across the QTL peak region (x-axis positions are megabases on the chromosome) Highlighted region is the QTL confidence interval.

**Figure 4:**
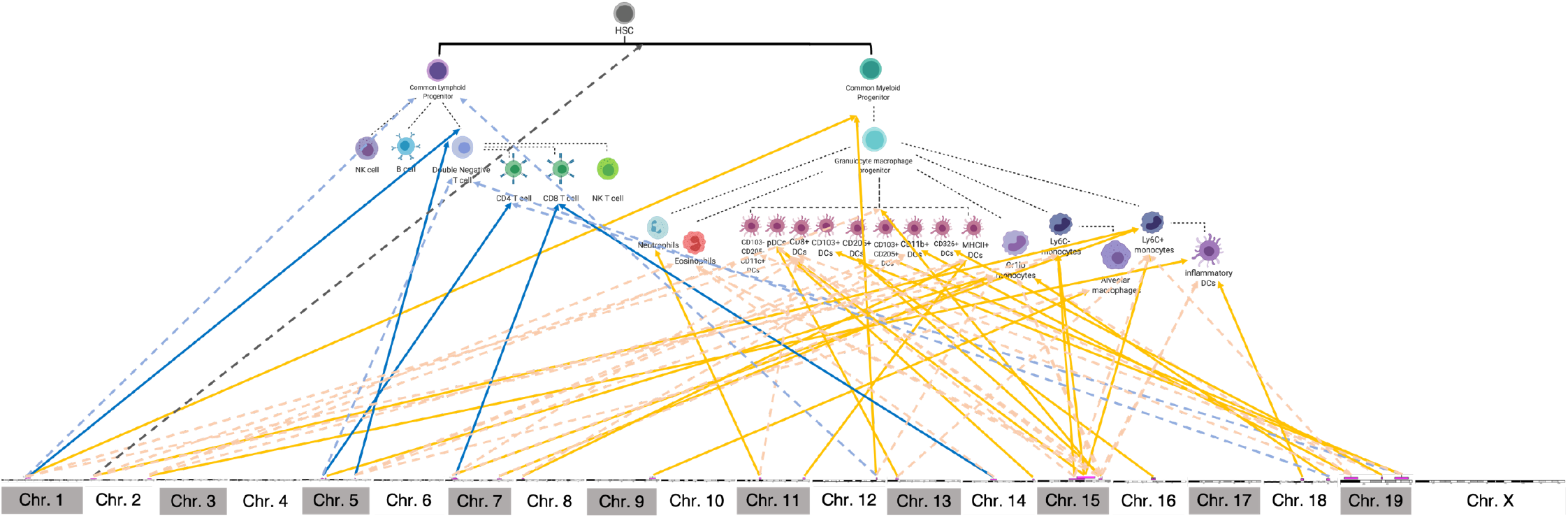
QTL identified for variation is lung immune cell populations at homeostasis in CC-F1 mice have broad direct and indirect effects. The mouse genome is represented linearly along the bottom with QTL represented as pink blocks (95% credible interval), while the upper panel is a developmental tree of the various immune populations we assessed. Blue solid arrows represent the direct effects for QTL and lymphoid populations they were mapped for, while blue dashed arrows indicate indirect effects between mapped QTL and lymphoid populations in the lungs of CC-F1 mice. Orange solid arrows represent the direct effects for QTL and myeloid populations they were mapped for, while orange dashed arrows indicate indirect effects between mapped QTL and myeloid populations in the lungs of CC-F1 mice. Grey solid arrows indicate indirect effects on both lymphoid and myeloid populations. Indirect effects were determined using our linear model analysis.

**Figure 5:**
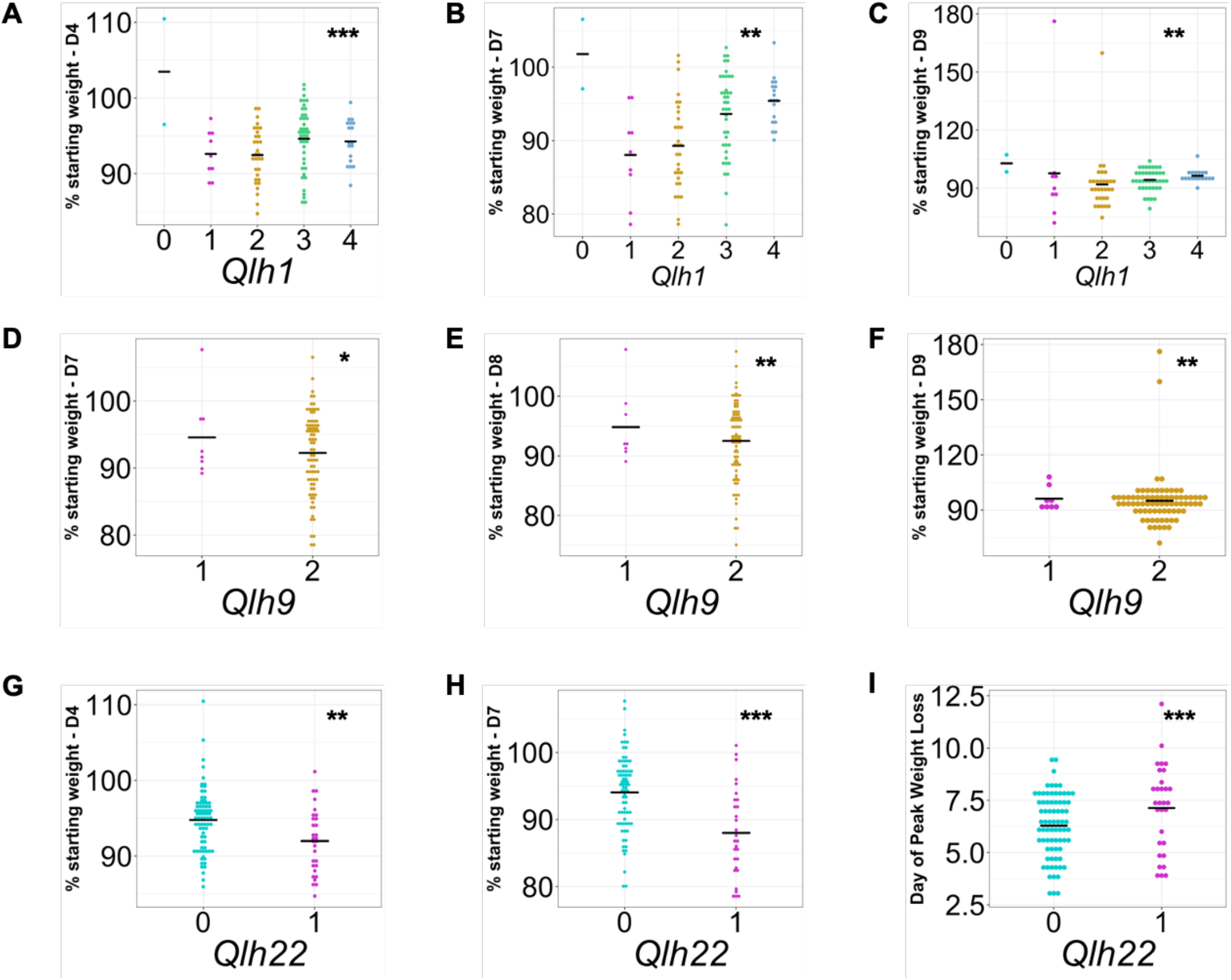
Relationships between QTL mapped for baseline immune cell frequencies in the lungs of CC-F1 mice and influenza A virus induced disease. Each point represents the mean value for each CC-F1 cross and the mean for each haplotype group on the x-axis is represented as a black bar. (A – C) *Qlh1* (CD4+ T cells, 0 = homozygous A/J vs. other haplotypes), (D – F) *Qlh9* (plasmacytoid DCs, 1 = heterozygous WSB vs 2 = homozygous other haplotypes), and (G – I) *Qlh22* (Gr1^lo^ monocytes/macrophages, 0 = other haplotypes vs. 1 = heterozygous NZO) show a range of relationships with IAV-induced disease, as measured by weight loss. (*p < 0.1, **p < 0.05, ***p < 0.01)

Of note, we identified *Qlh6,* a locus responsible for variation in alveolar macrophage frequencies in the lungs on Chromosome 9. This locus spans an approximately 11Mb region between 107.39Mb – 118.3Mb. At this locus, the NOD haplotype increases the frequency of alveolar macrophages in the lungs, relative to other haplotypes (Figure 3B, 3C). An assessment of the effect of *Qlh6* showed that it controlled 19.8% of the phenotypic variation and 37.3% of the heritability (Supplemental Table 2). We also identified *Qlh27* for variation in neutrophil frequencies in the lungs on Chromosome 11 spanning a 3Mb region, 36.20 – 39.54Mb. At this locus, the BL/6J, 129, and PWK haplotypes decrease the frequency of neutrophils in the lungs, relative to other haplotypes (Figure 3D, 3E). *Qlh27* explains 8.9% of the phenotypic variation in neutrophils and 14.2% of the heritability.

In several cases we identified more complex interactions. For example, various aspects of T cell abundances are governed by 5 identified loci *(Qlh1, Qlh2, Qlh3, Qlh4, Qlh5,* Supplemental table 2). These loci affected total T cells (*Qlh3* – Chr. 1:62.34 – 72.27Mb, PWK – low; *Qlh4* – Chr. 5:116.82 – 118.82Mb, NZO – high, CAST – low); CD4^+^ T cells (*Qlh1* – Chr. 5:42.06 – 48.35Mb, BL/6J, CAST, NZO – high, A/J – low) and CD8^+^ T cells (*Qlh2* – Chr. 14:47.66 – 58.58Mb, A/J – high, NZO – low; *Qlh5* – Chr. 7:18.02 – 25.26Mb, 129, CAST – low). The combined effects of these loci impacted 26.6% of total T cell variation, 20.8% of CD4^+^ T cell variation, and 43.5% of CD8^+^ T cell variation. In the other instance, loci such as *Qlh8* (Chr. 15:55.48 – 81.32Mb, WSB – high) was associated with Ly6C^hi^ monocyte frequency as well as pDC frequencies; albeit with somewhat discordant allele effects (Supplemental Table 2). All told, our results highlight an abundant and complex genetic regulation of baseline lung immune populations.

### Candidate gene analysis

Having mapped multiple QTL associated with variation in both single, as well as multi-population regulation of baseline leukocyte populations, we attempted to resolve a subset of loci to identify candidate causal variants. Therefore, we used our previously established analysis pipeline (Noll et al., 2020), where we filtered gene variants at each locus based on gene expression patterns and whether polymorphisms fit with haplotype effects. First, we filtered genes by expression in relevant tissues (e.g. lungs and spleen) at baseline. Next, we looked for genes with single nucleotide polymorphisms (SNPs) that were specific to the variant haplotype under the QTL. Lastly, we identified high-priority candidate genes based on the effect of the variant on protein sequence (i.e. non-synonymous or splice variant), as these have better defined functional consequences. Based on these criteria, we have compiled a list of candidate genes for the 10 QTL that reached our p < 0.1 significance threshold (Supplemental Table 3), with this approach reducing the list of candidates under a QTL from 21 – 388 genes to 1 – 201 genes.

In several cases, our candidate filtering process either reduced the locus to one or a few high priority candidate genes or suggested the mechanism by which the locus influences cell populations. Under *Qlh2* (CD8+ T cells, Chr.14: 47.66 – 58.58Mb), there are 388 genetic elements, and 130 are expressed in relevant tissues. We further refined this locus to 4 high priority candidate genes, *Vmn2r89, Ripk3, Parp4,* and *Ktn1* with either A/J or NZO specific non-synonymous variants. *Qlh6* (Alveolar macrophages, Chr. 9: 107.39 – 118.28Mb) contains 193 genetic elements, 112 of those are expressed in relevant tissues. We identified a single high priority candidate gene with NOD specific non-synonymous variants under *Qlh6*, *Pdcd6ip*, which codes for ALIX. Under *Qlh13 (*CD103^+^ DCs, CD103^+^CD205^+^ DCs, CD103^+^CD205^+^ DCs (prop. of Lin^-^), Chr. 19: 46.07 – 58.15Mb), there are 288 genetic elements, and 160 are expressed in relevant tissues. We identified 5 high priority candidate genes with NZO specific non-synonymous variants, *Loxl4, Pyroxd2, Hps1, Lzts2,* and *Trim8*, under *Qlh13*. In contrast to clear evidence of protein functional differences (missense and nonsense variants) gene regulation differences can be important in controlling biological processes, and several loci had candidate evidence suggesting gene regulation was important. *Qlh9* (pDCs, Chr. 14: 119.44 – 123.31Mb) there are 35 total genetic elements, with 20 expressed in relevant tissues. There are no genes under the locus that have WSB specific non-synonymous variants. However, there are 18 genes with WSB specific variants that do not change protein-coding, which suggests that the causal variant under this locus likely varies at the level of gene expression. Lastly, under *Qlh14* (Lineage-cells, Chr. 12: 106.34 – 107.94Mb) there are 21 genetic elements, and only one gene, *Rit1*, is expressed in a relevant tissue and contains WSB specific variants that do not change protein-coding.

### Relationships between baseline immune lung QTL and other homeostatic immune phenotypes in the lungs of CC-F1 hybrids

Our initial examination of the immune population structure (Figure 2) across our population suggested that there might be loci impacting the composition of multiple immune populations. In general, QTL tended to be identified for a single trait, despite some of the strong correlations noted. Alveolar macrophages and natural killer cells (ρ = 0.419) were highly correlated in the lungs, but we only identified a QTL for alveolar macrophages *(Qlh6,* Figure 3B), and no loci for NK cells (Supplemental Table 2). Even when both traits had QTL identified, such as inflammatory DCs (*Qlh23*, *Qlh24*) and MHCII+ DCs (*Qlh25*, *Qlh26*) they didn’t overlap despite strong correlations. Given the complex nature of genetic architecture across traits, we have previously used more simplified causal models to identify relationships between QTL and other phenotypes (Noll et al., 2020). We applied this approach across our lung immune cell population, asking whether the causal haplotype(s) under QTL identified for specific lung phenotypes (e.g. *QIh1*/CD4 T cells) were associated with the abundance of other cell populations. Our set of 28 loci were directly associated with abundance differences for 24 cell populations, leaving a possible 1478 additional associations which we could assess. We identified 205 additional relationships (p < 0.05 per test), with 55 of these remaining significant with an FDR correction applied (Supplemental Table 4) across these 28 loci. These effects were spread across the identified QTL (range 0 – 6, median 1 additional significantly affected populations per QTL; Supplemental Table 4). We found that the majority of loci (21/28) were associated with at least 1 additional phenotype, with a set of 5 of these loci associated with 4 or more additional phenotypes.

In the case of highly correlated populations such as MHCII^+^ DCs and inflammatory DCs (p=0.6179), our analysis did identify that *Qlh23* (inflammatory DCs, Chr. 2: 164.18 – 173.79Mb) was also associated with the frequency of MHCII^+^ DCs. Furthermore, this approach can also elucidate genetic regulation even when cellular phenotypes are not highly correlated in their overall abundances. For example, *Qlh2* (CD8^+^ T (proportion of total T cells)) is also associated with the frequency of eosinophils in the lungs (ρ = −0.20156). *Qlh27* (Neutrophils) is also associated with the frequency of CD8^+^ DCs in the lungs (ρ = −0.0692). Notably, *Qlh19* (CD11b^+^ DCs, Chr.19: 3.15 – 14.88Mb) is associated with four additional phenotypes including, Gr1lo monocytes/macrophages (ρ= 0.1211), Ly6C^hi^ monocytes/macrophages (ρ= −0.0395), CD103^-^CD205^-^DCs (ρ= 0.0323), and CD4^+^ T cells (proportion of total T cells) (ρ = −0.1206), suggesting that genes under loci such as *Qlh19*, may act as ‘master regulators’ of the homeostatic immune landscape in the lungs. This approach can elucidate co-QTL for both highly correlated cellular phenotypes, and cellular phenotypes with no phenotypic relationship.

### Relationships between baseline immune lung QTL and respiratory virus-induced phenotypes

Several immune cell populations in the lungs have been found to play vital roles in the response to respiratory virus infection (Channappanavar et al., 2016; Dalskov et al., 2020; LeMessurier et al., 2020). We therefore sought to determine if any of the genetic loci identified in our study showed an association with influenza A virus (IAV)-induced disease phenotypes. To do this, we used existing data from a similar panel of CC-F1 mice that were infected with IAV. Using all mapped QTL, we scored each type of CC-F1 hybrid based on their haplotype at the mapped loci. We then asked if there was any relationship between the haplotype score at the mapped locus and IAV-induced phenotypes. We found that four loci showed no relationship with gross IAV-induced disease, 13 loci showed one highly significant or one or more suggestive/significant associations with a measure of gross IAV-induced disease, and seven loci showed a highly significant association with three or more aspects of IAV-induced disease (Figure 6, Supplemental Table 5). Of note, *Qlh1* was identified with an A/J allele reducing CD4^+^ T cell abundance in the lungs, and CC-F1s with this A/J allele showed reduced weight loss at days 1-5 and days 7-9 post infection (Figure 6A – 6C). Additionally, *Qlh9* was identified with the WSB allele decreasing the frequency of pDCs in the lungs, and CC-F1s with the WSB allele at this locus showed reduced weight loss at days 7-9 post-infection (Figure 6D – 6F). Lastly, *Qlh22* was identified with a NZO allele increasing Gr1^lo^ monocyte/macrophage abundance in the lungs, and CC-F1s with the NZO allele at this locus showed enhanced weight loss at days 4-10 post-infection (Figure 6G – 6I), Supplemental Table 5).

## Discussion

Immune homeostasis is a critical mechanism through which the immune system balances its responsibility to protect against foreign insult and its potential to cause self-harm (Crimeen-Irwin et al., 2005). In contrast to more systemic immune homeostasis, the lungs represent a site that is regularly exposed to immune insults and as such presents an environment where maintenance of immune homeostasis in the face of frequent immune stimulation is critical. However, immune homeostasis, is a challenging phenotype to assess in humans due to differences in development, immunological history (Orrù et al., 2013), and tissue compartment access. As such, mouse models have been critical for understanding the genetic regulation of immune development and homeostasis (Falk et al., 1996; Graham et al., 2017; Kitamura et al., 1991; Krištić et al., 2018; Lansford et al., 1998; 2018b). We extend this work and describe an abundant and complex genetic regulation of multiple cell populations in the immune exposed lung. Importantly, we show that lymphocytic, inflammatory, resident (tissue unique) and antigen-presenting populations all show genetic regulation in this tissue and were able to identify genetic loci and candidate genes associated with 37 of these populations. We also demonstrated that the standard mouse strains BALB/cJ and BL/6J, which are heavily used to study lung immunology and the host response to pulmonary infection, are outliers for many immune phenotypes. Lastly, we demonstrate that genetic loci associated with variation in multiple aspects of pulmonary immune homeostasis can also be associated with variation in disease outcome following influenza infection, which suggests that genetic factors regulating baseline lung immune status have a significant impact on an individual’s subsequent susceptibility to respiratory infection. Altogether, these data highlight the critical importance of genetic variation on maintaining productive immune homeostasis, as well as the utility of understanding genetic regulation of lung immune homeostasis in the context of susceptibility to respiratory virus infection.

Experimental genetic mapping populations have classically been designed between pairs of inbred strains (e.g. F2 populations) to focus on specific phenotypes (Scalzo et al., 1995). In contrast, genetic reference populations (Mackay et al., 2012; Huang et al., 2015; Threadgill et al., 2012) have been designed to probe the role of genetic variation on traits in a *de novo* fashion. Our work here highlights the utility of using the latter approach. Namely, that by looking across a genetically diverse population, we were able to capture a greater breadth of genetically controlled immunological diversity. This manifests in our results showing that classic immunological models such as C57BL/6J and BALB/cJ are often phenotypically immune outliers, such as having a very low CD4/CD8 T-cell ratio, and highly abundant alveolar macrophage populations (Figure 1; similar outlier results being described in (Graham et al., 2017)). More than this, we also identified CC-F1s which showed other extreme immune homeostatic conditions. For example, two different CC-F1 hybrids, CC017xCC004 and CC005xCC001, showed eosinophilia (<30% of their lung immune compartment being eosinophils). Thus, exploring the larger genetic space afforded in a genetic reference panel uncovers a variety of new observations and potential models with regards to ‘normal’ immune homeostasis across inbred mouse strains.

By utilizing a large population of CC-F1 hybrids, we were able to further identify a number of QTL which: were distributed across the genome; affected a variety of immune phenotypes; and were driven by alleles from each of the eight founder strains of the CC. These results are in-line with our earlier observations of phenotypic diversity: that shuffling a variety of causal alleles across a population will lead to unique immune homeostatic phenotypes. The majority of the loci we identified impacted multiple cell populations. For example, *Qlh19* impacting Gr1^lo^ monocytes/macrophages, Ly6C^hi^ monocytes/macrophages, CD103^-^CD205^-^DCs, and CD4^+^ T cells; or *Qlh4* impacting CD103^+^, CD205^+^ DCs, CD11b^+^ DCs/interstitial macrophages (CD11b^+^ DCM), MHCII^+^ DCs, and alveolar macrophages in addition to the phenotypes they were initially mapped for. These broadly associated loci are suggestive of ‘master regulators’ of immune status. Some loci, such as *Qlh2* highlight potentially intriguing relationships. *Qlh2* was mapped due to its influence on the relative abundance of CD8^+^ T cells as a proportion of CD3^+^ T cells, and we further found that *Qlh2* was also associated with the proportion of eosinophils. To our knowledge, there have been no documented studies pointing to genes important in commonly regulating the abundance of these two cell populations, although there is significant evidence that there is crosstalk between eosinophils and CD8^+^ T cells, especially in pulmonary disease (Akuthota et al., 2008; Swain et al., 2006; Lee et al., 2001). Further studies assessing how these QTL directly impact the development, migration and tissue residency of each cell type, and whether these cell populations then cross-regulate each other is likely to provide important new insights into the regulation of (e.g. CD8^+^ T cells and eosinophils) homeostasis in the lungs.

A key goal of genetic mapping is not just to identify polymorphic regions, but the actual causal variants driving the phenotypic differences. Depending on the complexity of allele effects, and the underlying genetic variants (e.g. regulatory, structural), these candidate analyses can have differing levels of success (Ho et al., 1998; Idris et al., 1999; Noll et al., 2019; Yokoyama et al., 1997). For seven loci, we were able to reduce our loci to a small number of high priority candidate genes, with our analysis also identifying broader lists of candidate genes for the other QTL that reached our significance threshold (p < 0.1) (Supplemental Table 1), and we are in the process of further resolving candidate genes under these loci using our established methods (Noll et al., 2020). In the case of *Qlh6*, which is associated with variation in alveolar macrophages, we were able to resolve this locus down to a single high priority candidate gene, *Pdcd6ip*, which codes for the protein ALIX. ALIX is a member of the ESCRT pathway and is involved in extracellular vesicle (EV) transport (Fujita et al., 2018). EVs are known to be important for mediating crosstalk between epithelial cells and alveolar macrophages (Bissonnette et al., 2020). However, ALIX’s specific role in alveolar macrophage development or function has not been evaluated, and studies dissecting this impact would advance our understanding of alveolar macrophage biology. Another QTL with a priority candidate gene was *Qlh8*, mapped for Ly6C^hi^ monocytes, ratio of Ly6C^hi^ to Ly6C^-^monocytes, and pDCs. *Qlh8* includes the *Ly6* locus, pointing to putative effects in *Ly6c* itself driving these differences. However, the allele effects are slightly different for each phenotype. For the ratio of Ly6C^hi^ to Ly6C-phenotype, the CAST and WSB haplotypes are driving the QTL, while for the other phenotypes the only driver haplotype is WSB. There are no missense variants shared by CAST and WSB at this locus, although the CAST haplotype has missense variants in both *Lyc1* and *Ly6c2*. Given the complexity of allele effects, it is possible that functional differences in the CAST Ly6C heterodimer are driving some of the variation, while the variants in the WSB allele suggests a more complex interplay between gene regulation and function. In addition, *Qlh11*, mapped for Ly6C- monocytes, is proximal to the *Ly6c* locus, suggesting potential longer-distance regulatory differences affecting the Ly6C locus. In total, these complex allelic interactions point to a mixture of long- and short-range regulatory differences, combined with potential functional protein differences in interacting to drive monocyte homeostasis.

While we identified a great number of large effect QTL, and despite the high heritability of all cell population identified in this study, we were not able to identify QTL impacting each of these populations. For example, DN (CD4-CD8-CD3+) T cells are a population associated with autoimmune disorders (Alexander et al., 2020; Lohani et al., 2021). In our CC-F1 population, we found ~83% of the variation in their abundance attributable to genetic effects, yet we found no loci associated with this population. The power to map genetic effects is determined by many factors, including the number of loci impacting the disease, and also the gene-environment interactions impacting these cell types (Xue et al., 2020). Our study was designed to minimize environmental variation, such as nutrition, cage and housing effects. However, the contrast between PBS and no stimulation shows that there can be perturbation effects that could add noise and confound analysis. More likely however, is that many loci work together to determine cell population abundances in the lung. We observed this with eosinophils (50% heritable) in our population. We did not identify any QTL directly for eosinophil variation in our population, but using our simplified causal models identified 6 QTL for other immune cell populations in the lungs were significantly associated with eosinophil variation. As such, for many of our unmapped cell populations, or those with unexplained heritability, it is likely that multiple loci with smaller effects are driving these responses. Future studies wishing to understand the genetic regulation of these populations would be advised to use more targeted mapping crosses (Gu et al., 2020) designed to simplify the genetic architecture driving the regulation of these immune populations.

Normal immune homeostasis should allow for individuals to successfully respond to insults such as infections. We took advantage of our previously published data of matched CC-F1 mice infected with influenza (Maurizio and Ferris, 2017; Noll et al., 2020) to determine whether the genetic loci controlling immune homeostasis also were associated with influenza disease outcomes. Here we show that genetic regulation of homeostatic immune cell frequency in the lungs of CC-F1 mice may have subsequent impacts on influenza-induced disease. We found that approximately half of our identified loci had some association with IAV induced disease, demonstrating the importance of understanding homeostatic immunity to further interpret immune responses to respiratory virus infection. Importantly, some loci, such as *Qlh1* (decreased CD4^+^ T-cells) and *Qlh2* (increased CD8^+^ T-cells) showed protection from severe weight loss across the course of infection. These results are in line with the literature, which shows that CD8+ T cells are primarily responsible for clearing influenza A virus infection (Fiege et al., 2019), and extend these findings to suggest that even prior to influenza specific adaptive CD8^+^ T cell responses, residency of CD8+ T cells in the lungs before infection might play a key role in the response to infection. Similarly, we identified loci increasing Ly6C^hi^ monocyte abundance (*Qlh7*) and increasing alveolar macrophage abundance which were significantly associated with improved influenza disease responses (Supplemental Table 5).

It is highly likely that the diverse nature of the genetic control of lung immune populations may lead to differences in infection responses in a pathogen-specific manner. For example, studies of Severe Acute Respiratory Syndrome coronavirus (SARS-CoV) have shown that alveolar macrophages may contribute to disease severity by dampening dendritic and CD8^+^ T cell activation (Zhao et al., 2009), while Ly6C^hi^ inflammatory monocytes promote SARS-CoV-induced pulmonary damage and disease (Channappanavar et al., 2016). These results stand in contrast to the observed associations with IAV disease outcome we identified above. Such results highlight the complex nature of immune-pathogen interactions, and also point to a larger utility in studying infectious responses across a range of models. This is most relevant in the context of (e.g) the current SARS-CoV-2 pandemic, or other endemic respiratory insults (e.g. asthma) where assessing responses across sets of mouse strains which vary in different components of their immune homeostasis (e.g. pro- vs anti- inflammatory environments; CD4 to CD8 T cell ratios) might help clarify correlates of protection or severe disease.

A large and growing body of literature exists describing the role of immune homeostasis on phenotypes related to vaccination, infection and aberrant immunity (Crimeen-Irwin et al., 2005; Gnjatic et al., 2017; Graham et al., 2017; HIPC-CHI Signatures Project TeamHIPC-I Consortium, 2017; Tsang et al., 2014). Building on classic mouse models of immunity, we show that genetic variants drive immune homeostasis in both tissue- and cell type- specific manners. We also show that this regulation plays a role in viral disease responses. Writ large, we highlight the relevance of mouse GRPs in describing and characterizing immune regulation at basal states, as well as during infectious responses. Follow-up studies investigating the loci presented here and mechanistic studies of candidate genes underlying those loci could enhance our understanding of immune function in the lungs and the factors controlling lung specific makeup of various cell compartments at the steady state. Additionally, understanding lung immune function at the steady state will be important for investigating immune dysfunction in the context of allergy and autoimmunity, as well as immune responses to respiratory viral and bacterial pathogens.

## Methods

### Mice

Mice from 64 CC strains were obtained from the Systems Genetics Core Facility at the University of North Carolina-Chapel Hill (UNC) between July 2021 and July 2016. CC-F1 hybrids were generated by breeding the CC mice in our laboratory. Approximately 300 female mice were generated from 94 CC-F1 crosses of these 64 CC strains. At 4-6 weeks of age, mice were transferred from their breeding facility to a BSL-2 facility at UNC Chapel Hill. BL/6J and BALB/cJ mice were purchased from The Jackson Laboratory at eight weeks of age. Additional inbred CC mice from 6 CC strains were purchased from the Systems Genetics Core Facility at the University of North Carolina-Chapel Hill (UNC) in 2018. All mice were housed under specific-pathogen free conditions with a 12hr light/dark cycle, and food and water ad libitum in both UNC Chapel Hill vivaria described above. All experiments were conducted under compliance with IACUC protocols at UNC-Chapel Hill.

### Treatment and Cell preparation

8-12-week-old animals were anesthetized via Isoflurane inhalation and intranasally inoculated with 50uL of PBS only. 4 days later, animals were euthanized with isoflurane overdose and cardiac puncture. Mice were perfused with 10mL PBS and whole lungs were harvested. Lungs were digested in digest media (RPMI + 10% FBS + DNase I + Collagenase) at 37ºC for 90 minutes. Lung homogenates were treated with ammonium chloride potassium (ACK) lysis buffer to remove red blood cells, washed, filtered through 70um mesh, and resuspended in fluorescence-activated cell sorting (FACS) buffer (HBSS + 1% FBS + 0.01% sodium azide). Cell numbers were determined using a hemocytometer and trypan blue exclusion.

### Flow cytometry

Cells were plated at 1×10^7^ per mL in FACS buffer in 96-well polypropylene round-bottom plates. Cells were centrifuged at 1000 rpm for 4 minutes, resuspended in 100uL of fluorochrome-conjugated antibody dilution, and stained for surface markers for 45 minutes at 4ºC. The cells were washed 2x in FACS buffer, resuspended in 200 μL of FACS buffer and added to another 200 μL of fresh 4% paraformaldehyde (ACROS Organics, Thermo Fisher Scientific) in PBS and stored at 4°C in the dark until analysis on the flow cytometer. A Dako-Cytomation CyAn was used for all analysis and the data analyzed and recorded using the Summit software. The following antibodies were used: Ly6C-FITC (AL-21), SiglecF-PE (E50-2440), CD11c-PE Texas Red (N418), CD45R/B220-PerCP (RA3-6B2), Gr-1-PE-Cy7 (RB6-8C5), CD11b-eF450 (M1/70), CD45/LCA-APC (30-F11), MHCII-APC-eF780 (M5/114.15.2), CD94-FITC (18d3), CD3ε-PE (145-2C11), CD4-PE Texas Red (GK1.5), CD8α-PerCP (53-6.7), CD49b-PE-Cy7 (DX5), CD45/LCA-eF450 (30-F11), CD19-APC (6D5), CD45R/B220-APC-eF780 (RA3-6B2), CD3ε-FITC (145-2C11), CD45R/B220-FITC (RA3-6B2), CD19-FITC (6D5), CD103-PE Texas Red (2E7), CD326-PE-Cy7 (G8.8), CD205-APC (NLDC-145), CD45/LCA-APC-eF780 (30-F11). Flow cytometric analysis was completed using FlowJo v10 software (Tree Star, Ashland, OR). Debris, doublet events, auto-fluorescent cells, and CD45-events were eliminated from all samples prior to downstream cell population quantifications. Gating schemes used for flow cytometry analysis are shown in Supplemental Figures 1-3.

### Data processing

Raw event count values from flow cytometry gating were used to calculate proportions and scaled. Using the MASS package in R (version 3.5.1), Box-Cox transformation values were determined independently for each cell phenotypes and the phenotype values were transformed accordingly to follow a normal distribution. For all phenotypes, the average value was calculated for each CC-F1 line, which was used for QTL mapping.

### Heritability estimates

Heritability calculation were performed as described previously (Noll et al., 2020). Briefly, box-cox transformed phenotype values were used to fit a linear fixed-effect model. The coefficient of genetic determination was calculated as such:

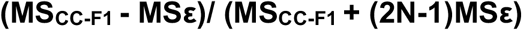

Where MS_CC-F1_ is the mean square of the CC-F1 and MSε is the mean square of the error using a N=3 as an average group size, as a measure of broad-sense heritability.

### QTL mapping

QTL mapping was performed as previously described (Noll et al., 2020). Briefly, we used the DOQTL (Gatti et al., 2014) package in the R statistical environment (version 3.5.1). A multiple regression is performed at each marker, assessing the relationship between the phenotype and the haplotype probabilities for each strain. LOD scores are calculated based on the increase in statistical fit compared to a null model, considering only covariates and kinship. To calculate significance thresholds, permutation tests were used to shuffle genotypes and phenotypes without replacement. We determined the 80^th^, 90^th^, and 95^th^ percentiles after 500 permutations as cutoffs for suggestive (both p<0.2 and p<0.1) and genome-wide significant (p<0.05. QTL intervals were determined using a 1.5 LOD drop.

### Phenotype correlations with mapped QTL

Correlations between identified QTL and other homeostatic and IAV infection-induced phenotypes were determine by comparing the goodness of model fit of mixed effect linear models using a partial fit F-test. CC-F1 haplotype scores were determined as the additive score of the dam and sire founder haplotype at the QTL peak marker. In both the base and full model, CC-F1 is a random effect variable and QTL haplotype score is a fixed effect variable. The full model tests whether including information about the haplotype at the mapped QTL explains more of the phenotypic variation than the CC-F1 cross alone.

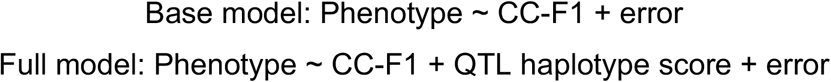

To assess the relationship between QTL and IAV-induced disease phenotypes, a *Mx1* haplotype score was included in the base and full model as a fixed effect variable. We have shown that variation in *Mx1*, a powerful host antiviral gene, explains the vast differences in disease susceptibility to IAV infection in the CC (Maurizio et al., 2018).

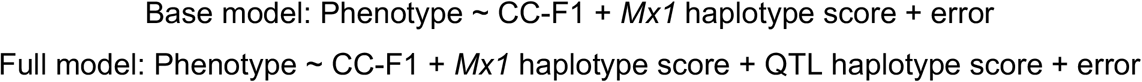

### Candidate gene refinement

Genetic elements under the locus were first narrowed by selecting those that were expressed in the lung and/or spleen at homeostasis. Expressed genes were then narrowed by selecting those that contain QTL haplotype-specific variants from the Sanger SNP database (Keane et al., 2011; Yalcin et al., 2011). These genes are listed as candidate genes under each QTL. High priority candidate genes are further defined as expressed genes that contain QTL haplotype-specific variants that are predicted to alter protein-coding (e.g. missense or nonsense variants).

