## Supplemental Data for "Genetic regulation of homeostatic immune architecture in the lungs of Collaborative Cross mice"

Supplemental Table 1: Effects of PBS-instillation on lung leukocyte populations in CC strains. Bolded rows indicate phenotypes where PBS treatment had a significant impact on cellular frequency across strains.

| PHENOTYPE | PHENOTYPIC DISTRIBUTION (untreated) | PHENOTYPIC DISTRIBUTION (PBS instillation) | PBS Treatment Effect p-value |
|---|---|---|---|
| $CD3^{+}$ T cells (proportion of $LCA^{+}$) | 0.249 – 0.921 | 0.206 – 0.711 | p = 0.4737 |
| **$CD4^{+}$ T cells (proportion of $LCA^{+}$)** | **0.001 – 0.132** | **0.008 - 0.107** | **p = 0.0177** |
| **$CD4^{+}$ T cells (proportion of $CD3^{+}$ T cells)** | **0.001 – 0.304** | **0.011 - 0.260** | **p = 0.0005** |
| DN T cells (proportion of LCA+) | 0.002 – 0.062 | 0 – 0.068 | p = 0.3433 |
| DN T cells (proportion of CD3+ T cells) | 0.005 – 0.067 | 0.001 – 0.099 | p = 0.2217 |
| $CD8^{+}$ T cells (proportion of $LCA^{+}$) | 0.087 – 0.795 | 0.125 – 0.569 | p = 0.2331 |
| **$CD8^{+}$ T cells (proportion of $CD3^{+}$ T cells)** | **0.323 – 0.863** | **0.349 – 0.801** | **p = 0.0093** |
| **CD4 -to- CD8 T cell ratio** | **0 – 0.470** | **0.010 – 0.430** | **p = 0.0012** |
| B cells (proportion of $LCA^{+}$) | 0.100 – 0.400 | 0.120 – 0.460 | p = 0.4471 |
| $Ly6C^{-}$ monocytes (proportion of $LCA^{+}$) | 0.002 – 0.036 | 0.002 – 0.011 | p = 0.5661 |
| $Ly6C^{-}$ monocytes (fraction of $Ly6C^{+/lo/-}$) | 0.090 – 0.600 | 0.090 – 0.470 | p = 0.4520 |
| $Ly6C^{+}$ monocytes (proportion of $LCA^{+}$) | 0.001 – 0.017 | 0.001 – 0.019 | p = 0.0882 |
| $Ly6C^{+}$ monocytes (fraction of $Ly6C^{+/lo/-}$) | 0.050 – 0.500 | 0.050 - 0.540 | p = 0.5805 |
| $Ly6C^{lo}$ monocytes (proportion of $LCA^{+}$) | 0.008 - 0.020 | 0.007 – 0.023 | p = 0.1247 |
| $Ly6C^{lo}$ monocytes (fraction of $Ly6C^{+/lo/-}$) | 0.310 – 0.650 | 0.360 – 0.660 | p = 0.7230 |
| Eosinophils (proportion of $LCA^{+}$) | 0.013 – 0.068 | 0.015 – 0.081 | p = 0.8329 |
| Neutrophils (proportion of $LCA^{+}$) | 0 – 0.036 | 0 – 0.016 | p = 0.5980 |
| Alveolar Macrophages (proportion of $LCA^{+}$) | 0.025 – 0.255 | 0.011 – 0.271 | p = 0.8321 |

| | | | |
|---|---|---|---|
| plasmacytoid DCs (proportion of $LCA^+$) | 0.037 – 0.241 | 0.035 – 0.217 | $p = 0.4348$ |

Supplemental Table 2: Summary of all QTL mapped ($p < 0.2$) and allele effects.

| NAME | PHENOTYPE | QTL INTERVAL | HAPLOTYPE EFFECTS | P-VALUE | HERITABILITY | PHENOTYPIC VARIATION EXPLAINED | GENETIC VARIATION EXPLAINED |
|---|---|---|---|---|---|---|---|
| *Qlh1* | **CD4+ T cells** (proportion of LCA+ cells) | Chr. 5:42.06 – 48.35Mb | BL/6J, CAST, NZO – high<br>A/J – low | 0.081 | 0.868 | 20.8% | 29.7% |
| *Qlh2* | **CD8+ T cells** (proportion of CD3+ T cells) | Chr. 14: 47.66 – 58.58Mb | A/J – high<br>NZO– low | 0.080 | 0.749 | 23.8% | 37.3% |
| *Qlh3* | **$CD3^+$ T cells** | Chr. 1:62.34 - 72.27Mb | PWK – low | 0.123 | 0.582 | 7.7% | 10.6% |
| *Qlh4* | **$CD3^+$ T cells** | Chr. 5: 116.82 – 118.82Mb | NZO – high<br>CAST – low | 0.179 | 0.582 | 18.9% | 31.9% |
| *Qlh5* | **$CD8^+$ T cells** (proportion of $LCA^+$ cells) | Chr. 7: 18.02 – 25.26Mb | 129, CAST – low | 0.124 | 0.737 | 19.7% | 27.2% |
| *Qlh6* | **Alveolar Macrophages** | Chr. 9: 107.39 – 118.28Mb | NOD – high | 0.025 | 0.536 | 19.8% | 32.6% |
| *Qlh7* | **$Ly6C^+$ monocytes** (proportion of $LCA^+$) | Chr. 2: 17.60 – 33.25Mb | WSB – high | 0.026 | 0.566 | 26.9% | 61.9% |
| *Qlh8* | **$Ly6C^+$ monocytes** (proportion of $LCA^+$)<br>**$Ly6C^+$ monocytes** (proportion of $Ly6C^{+/lo/-}$)<br>**plasmacytoid DCs**<br>**$Ly6C^-$ monocytes** (proportion of $Ly6C^{+/lo/-}$) | Chr. 15: 55.48 – 81.32Mb | WSB – high, CAST – low<br>WSB – high<br>WSB – high<br>CAST – high, WSB – low | 0.017<br>0.002<br>0.065<br>0.001 | 0.566<br>0.497<br>0.600<br>0.739 | 28.9%<br>34.1%<br>24.2%<br>32.5% | 55.4%<br>79.7%<br>41.2%<br>59.4% |
| *Qlh9* | **plasmacytoid DCs** | Chr. 14: 119.44 – 123.31Mb | WSB– low | 0.084 | 0.600 | 3.3% | 3.9% |
| *Qlh10* | **plasmacytoid DCs** | Chr. 13: 12.53 – 17.43Mb | BL/6J – high | 0.154 | 0.600 | 17.2% | 25.7% |
| *Qlh11* | **$Ly6C^-$ monocytes** | Chr. 15: 44.12 – 66.35Mb | CAST, NZO, NOD, A/J - high | 0.014 | 0.666 | 25.7% | 46.4% |
| *Qlh12* | **$CD11b^-$ cells**<br>**$CD11c^+$ cells** | Chr. 15: 86.88 – 91.33Mb | CAST – low | 0.086<br>0.081 | 0.565<br>0.526 | 3.2%<br>4.3% | 4.6%<br>6.3% |
| *Qlh13* | **$CD103^+$ DCs**<br>**$CD103^+$, $CD205^+$ DCs**<br>**$CD103^+$, $CD205^+$ DCs** (proportion of $Lin^-$) | Chr. 19: 46.07 – 58.15Mb | CAST – high, NZO – low<br>NZO – low<br>NZO – low | 0.193<br>0.131<br>0.086 | 0.539<br>0.372<br>0.358 | 0.51%<br>2.5%<br>0.32% | 1.2%<br>5.3%<br>0.63% |
| *Qlh14* | **$Lineage^-$ ($Lin^-$)** | Chr. 12: 106.34 – 107.94Mb | WSB – low | 0.085 | 0.782 | 10.7% | 14.1% |
| *Qlh15* | **$Lineage^-$ ($Lin^-$)** | Chr. 1: 59.24 – 62.47Mb | PWK – low | 0.124 | 0.782 | 2.1% | 2.6% |
| *Qlh16* | **$Ly6C^{lo}$ monocytes** (proportion of $Ly6C^{+/lo/-}$) | Chr. 7: 12.89 – 28.04Mb | NOD, 129 – high<br>BL/6J, CAST – low | 0.101 | 0.508 | 23.2% | 41.4% |
| *Qlh17* | **$CD205^+$ DCs** | Chr. 15: 90.61 – 91.33Mb | CAST – low | 0.163 | 0.627 | 2.5% | 3.4% |
| *Qlh18* | **$CD205^+$ DCs** | Chr. 16: 50.90 – 57.51Mb | NZO, A/J – high<br>CAST, PWK – low | 0.179 | 0.627 | 9.3% | 13.5% |
| *Qlh19* | **$CD11b^+$ DCs** | Chr.19: 3.15 – 14.88Mb | A/J – low | 0.138 | 0.499 | 7.2% | 11.9% |
| *Qlh20* | **$Gr1^{lo}$ monocytes/macrophages** | Chr. 18: 78.23 – 82.03Mb | NZO – low | 0.119 | 0.531 | 1.9% | 2.4% |
| *Qlh21* | **$Gr1^{lo}$ monocytes/macrophages** | Chr. 5: 50.56 – 53.57Mb | BL/6J – high | 0.197 | 0.531 | 4.8% | 12.7% |

| | | | | | | | |
|---|---|---|---|---|---|---|---|
| *Qlh22* | **Gr1$^{lo}$ monocytes/macrophages** | Chr. 7: 115.33 – 126.25Mb | NZO – high | 0.201 | 0.531 | 2.0% | 3.0% |
| *Qlh23* | **Inflammatory DCs** | Chr. 2: 164.18 – 173.79Mb | WSB – high<br>NZO, BL/6J – low | 0.132 | 0.654 | 20.4% | 29.8% |
| *Qlh24* | **Inflammatory DCs** | Chr. 18: 45.68 – 49.46Mb | PWK – low | 0.168 | 0.654 | 5.3% | 6.3% |
| *Qlh25* | **MHCII$^{+}$ DCs** | Chr. 8: 10.94 – 15.01Mb | PWK, WSB – high<br>129 – low | 0.159 | 0.728 | 16.2% | 21.9% |
| *Qlh26* | **MHCII$^{+}$ DCs** | Chr. 11: 110.16 – 113.59Mb | NZO – high<br>BL/6J – low | 0.192 | 0.728 | 21.9% | 31.2% |
| *Qlh27* | **Neutrophils** | Chr. 11: 36.20 – 39.54Mb | 129, BL/6J, PWK – low | 0.171 | 0.478 | 8.9% | 14.2% |
| *Qlh28* | **CD326$^{+}$ DCs** | Chr. 19: 34.69 – 36.13Mb | CAST - high | 0.179 | 0.612 | 8.6% | 11.1% |

Supplemental Table 3: Candidate gene lists for highly significant and significant QTL ($p < 0.1$).

| NAME | PHENOTYPE | QTL INTERVAL | HAPLOTYPE EFFECTS | CANDIDATE GENES | HIGH PRIORITY CANDIDATES |
|---|---|---|---|---|---|
| *Qlh1* | **CD4+ T cells** (proportion of LCA+ cells) | Chr. 5:42.06 – 48.35Mb | BL/6J, CAST, NZO – high<br>A/J – low | *Cpeb2, C1qtnf7, Cc2da, Fbxl5, Bst1, Cd38, Fgfbp1, prom1, Tapt1, Ldb2, Qdpr, Lap3, Med28, Fam184b, Ncapg, Lcorl, Slit2* | *Cpeb2, Cc2da, Prom1, Fam184b, Ncapg, Slit2* |
| *Qlh2* | **CD8+ T cells** (proportion of CD3+ T cells) | Chr. 14: 47.66 – 58.58Mb | A/J – high<br>NZO – low | *Ktn1, Peli2, Tmem260, Exoc5, Ap5m1, Slc35f4, Olfr736, Olfr747,Tlr11, Ttc5, Ccnb1ip1, Parp2, Tep1, Osgep, Pnp, Ang, Pnp2, Rnase4, Rnase6, Vmn2r89, Mettl17, AY358078, Hnrnpc, Arhgef40, Zfp219, Rpgrip1, Supt16, Chd8, Rab2b, Mettle3, Slc7a7, Sall2, Dad1, Oxa1l, Rem2, Haus4, Ajuba, 4931414P19Rik, Acin1, Slc7a8, Homez, Ppp1r3e, Pabpn1, Jph4, Carmil3, Tm9sf1, Nedd8, Tinf2, Gm5801, Ngdn, Thtpa, Zfhx2, Dhrs4, Nrl, Rec8, Tgm1, Ripk3, Rnf17, Parp4, Mphosph8, Pspc1, Gjb2, Zmym2, Il17d, Eef1akmt1, Xpo4, Lats2, Zdhhc20, Micu2* | *Vmn2r89, Ripk3, Parp4, Ktn1* |
| *Qlh6* | **Alveolar Macrophages** | Chr. 9: 107.39 – 118.28Mb | NOD – high | *Cacnad2d, Sema3b, Rbm5, Rbm6, Mon1a, Mst1r, Gmppb, Rnf123, Lamb2, Qars, Slc26a6, Col7a1, Pfkfb4, Shisa5, Fbxw22, Camp, Map4, Smarcc1, Ptpn23, Klhl18, Kif9, Setd2, Pth1r, Myl3, Als2cl, Tdgf1, Lrrfip2, Epm2aip1, Arpp21, Pdcd6ip, Clasp2, Glb1, Trim71, Cnot10, Cmtm8, Gpd1l, Osbpl10, Stt3b, Gadl1, Tgfbr2, Rbms3, Azi2* | *Pdcd6ip* |
| *Qlh7* | **Ly6C$^{+}$ monocytes** (proportion of LCA$^{+}$) | Chr. 2: 17.60 – 33.25Mb | WSB – high | *Nebl, Skida1, Mllt10, Dnajc1, Commd3, Bmi1, Spag6, Pip4k2a, Armc3, 4921504E06Rik, Otud1, Etl4, Arhgap21, Enkur, Thnsl1, Gad2, Apbb1ip, Pdss1, Abi1, Acbd5, Mastl, Yme1l1, Spopl, Hnmt, Il1f9, Il1rn, Psd4, Cacna1b, Ehmt1, Arrdc1, Zmynd19, Dph7, Pnpla7, Nsmf, Entpd8, Noxa1, Tor4a, Nelfb, Tubb4b, Uap1l1, Fut7, Abca2, Clic3, Paxx, Ptgds, Lcn12, C8g, Fbxw5, Traf2, Mamdc4, Rabl6, Tmem141, Kcnt1, Ubac1, Nacc2, Qsox2, Ccdc187, Gpsm1, Snapc4, Inpp5e, Sec16a, Notch1, Egfl7, Lcn4, Surf1, Surf4, Rexo4, Adamts13, Cacfd1, Slc2a6, Mymk, Adamtsl2, Fam163b, Sardh, Vav2, Brd3, Wdr5, Rxra, Col5a1, Fcnb, Ppp1r26, 1700007K13Rik, Mrps2, Gbgt1, Gtf3c5, Gfi1b, Spaca9, Ak8, Gtf3c4, Ddx31, Cfap77, Ttf1, Setx, Ntng2, Med27, Rapgef1, Slc27a4, Urm1, Cercam, Odf2, Gle1, Wdr34, Set, Pkn3, Zer1, Tbc1d13, Endog, Spout1, Kyat1, Lrrc8a, Phyhd1, Sh3glb2, Miga2, Ptpa, Ier5l, Cstad, 1700001O22Rik, Ptges, Tor1b,* | *Nsmf, Lcn12, Mamdc4, Ccdc187, Lamc3, Lrsam1* |

| | | | | | |
|---|---|---|---|---|---|
| | | | | *BC005624, Usp20, Fnbp1, Gpr107, Ncs1, Ass1, Fubp3, Exosc2, Abl1, Fibcd1, Lamc3, Aif1l, Nup214, Fam78a, Plpp7, Prrc2b, Pomt1, Swi5, Golga2, Dnm1, Ciz1, Ptges2, Slc25a25, Naif1, Fam102a, St6galnac4, St6galnac6, Ak1, Eng, Fpgs, Cdk9, Sh2d3c, Tor2a, Ttc16, Cfap157, Stxbp1, Fam129b, Lrsam1, Rpl12, Slc2a8, Garnl3, Ralgps1* | |
| *Qlh8* | **Ly6C$^+$ monocytes** (proportion of LCA$^+$)<br>**Ly6C$^+$ monocytes** (proportion of Ly6C$^{+/lo/-}$)<br>**plasmacytoid DCs**<br>**Ly6C$^-$ monocytes** (proportion of Ly6C$^{+/lo/-}$) | Chr. 15: 55.48 – 81.32Mb | WSB – high, CAST – low<br>WSB – high<br>WSB – high<br>CAST – high, WSB – low | *Col14a1, Mrpl13, Mtbp, Sntb1, Has2, Slc22a22, Zhx2, Derl1, Tbc1d31, Fam83a, 9130401M01Rik, Zhx1, Atad2, Wdyhv1, Fbxo32, Fam91a1, Tmem65, Trmt12, Rnf139, Tatdn1, Ndufb9, Mtss1, Sqle, Washc5, Nsmce2, Trib1, Myc, Gsdmc, Gsdmcl-ps, Fam49b, Asap1, Adcy8, Efr3a, Hhla1, Lrrc6, Tmem71, Phf20l1, Tg, Sla, Ndrg1, St3gal1, Zfat, Khdrbs3, Fam135b, Col22a1, Trappc9, Chrac1, Ago2, Ptk2, Dennd3, Slc45a4, Gpr20, Ptp4a3, Mroh5, Adgrb1, Arc, Jrk, Them6, Slurp1, Lypd2, Slurp2, Lynx1, Ly6d, Ly6k, Gml2, Ly6e, Ly6i, Ly6a, Ly6c1, Ly6c2, Ly6f, Ly6h, Gpihbp1, Zfp41, Top1mt, Rhpn1, Mafa, Zc3h3, Eef1d, Gsdmd, Naprt, Pycrl, Tsta3, Zfp623, Zfp707, Ccdc166, Mapk15, Fam83h, Scrib, Puf60, Plec, Parp10, Grina, Oplah, Exosc4, Gpaa1, Cyc1, Hgh1, Mroh1, Scx, Bop1, Hsf1, Dgat1, Fbxl6, Adck5, Cpsf1, Slc39a4, Tonsl, Cyhr1, Foxh1, Ppp1r16a, Gpt, Lrrc14, Lrrc24, Arhgap39, Zfp251, Zfp7, Commd5, Zfp647, 1110038F14Rik, Mb, Apol6, Rbfox2, Apol7a, Apol9a, Apol7b, Apol10a, Apol7c, Apol10b, Apol11b, Apol7e, Apol9b, Myh9, Txn2, Foxred2, Eif3d, Ift27, Pvalb, Ncf4, Csf2rb2, Csf2rb, Tst, Mpst, Tmprss6, Il2rb, C1qtnf6, Sstr3, Rac2, Cyth4, Elfn2, Mfng, Card10, Cdc42ep1, Lgals2, Gga1, Sh3bp1, Lgals1, Nol12, Triobp, H1f0, Gcat, Ankrd54, Eif3l, Micall1, 1700088E04Rik, Polr2f, Sox10, Pick1, Slc16a8, Baiap2l2, Pla2g6, Maff, Tmem184b, Csnkle, Kdelr3, Ddx17, Dmc1, Fam227a, Tomm22, Josd1, Gtpbp1, Sun2, Dnal4, Cbx6, Apobec3, Cbx7, Pdgfb, Rpl3, Syngr1, Tab1, Mgat3, Mief1, Rps19bp1, Cacna1i, Grap2, Fam83f, Tnrc6b, Adsl, Sgsm3, Mchr1, Slc25a17* | *Mtbp, Slc22a22, Zhx2, Fam83a, Zhx1, Fam91a1, Trmt12, Myc, Gsdmc, Adcy8, Hhla1, Lrrc6, Tmem71, Tg, Zfat, Fam135b, Col22a1, Trappc9, Dennd3, Mroh5, Ly6e, Ly6c1, Ly6c2, Top1mt, Zc3h3, Gsdmd, Zfp707, Ccdc166, Scrib, Plec, Parp10, Mroh1, Gpaa1, Fbxl6, Cpsf1, Slc39a4, Ppp1r16a, Zfp251, Zfp7, 1110038F14Rik, Apol9a, Apol7c, Apol11b, Apol7e, Apol9b, Ncf4, Csf2rb2, Mpst, Tmprss6, Il2rb, Sstr3, Card10, Nol12, Triobp, Gcat, Micall1, Fam227a, Gtpbp1, Sun2, Cbx7, Rpl3, Cacna1i , Syngr1, Tab1, Mgat3, Rps19bp1, Adsl* |
| *Qlh9* | **plasmacytoid DCs** | Chr. 14: 119.44 – 123.31Mb | WSB – low | *Hs6st3, Oxgr1, Mbnl2, Rap2a, Ipo5, Farp1, Stk24, Slc15a1, Dock9, Ubac2, Timm8a2, Tm9sf2, Clybl, Gm5089, Pcca, Ggact, Tmtc4, Nalcn* | |

| | | | | | |
|---|---|---|---|---|---|
| *Qlh11* | **Ly6C⁻ monocytes** | Chr. 15: 44.12 – 66.35Mb | CAST, NZO, NOD, A/J - high | *Trhr, Nudcd1, Eny2, Pkhd1l1, Ebag9, Sybu, Csmd3, Trps1, Eif3h, Utp23, Rad21, Aard, Med30, Ext1, Samd12, Tnfrsf11b, Colec10, Mal2, Enpp2, Taf2, Dscc1, Deptor, Col14a1, Mrpl13, Mtbp, Sntb1, Has2, Slc22a22, Zhx2, Derl1, Tbc1d31, Fam83a, 9130401M01Rik, Zhx1, Atad2, Wdyhv1, Fbxo32, Fam91a1, Tmem65, Trmt12, Rnf139, Tatdn1, Ndufb9, Mtss1, Sqle, Washc5, Nsmce2, Trib1, Myc, Gsdmc, Gsdmcl-ps, Fam49b, Asap1, Adcy8, Efr3a, Hhla1* | *Taf2, Hhla1, Atad2, Sybu, Pkhd1l1, Csmd3, Trps1, Tnfrsf11b, Colec10, Enpp2, Deptor, Col14a1, Mtbp, Slc22a22, Zhx2, Fam83a, Zhx1, Fam91a1, Trmt12, Myc, Gsdmd, Adcy8* |
| *Qlh12* | **CD11b⁻ cells**<br>**CD11c⁺ cells** | Chr. 15: 86.88 – 91.33Mb | CAST – low | *Brd1, Zbed4, Alg12, Creld2, Pim3, Ttll8, Mlc1, Mov10l1, Trabd, Tubgcp6, Mapk12, Mapk11, Plxnb2, Dennd6b, Ppp6r2, Sbf1, Adm2, Lmf2, Klhdc7b, Chkb, Mapk8ip2, Shank3, Acr, Rabl2, Syt10, Deaf1, Cpne8, Kif21a, Abcd2, Slc2a13* | *Tubgcp6, Adm2, Acr, Kif21a, Abcd2* |
| *Qlh13* | **CD103⁺ DCs**<br>**CD103⁺, CD205⁺ DCs**<br>**CD103⁺, CD205⁺ DCs**<br>(proportion of Lin⁻) | Chr. 19: 46.07 – 58.15Mb | CAST – high, NZO – low<br>NZO – low<br>NZO – low | *Plce1, Noc3l, Hells, Cyp2c69, Cyp2c54, Pdlim1, Sorbs1, Tctn3, Entpd1, Ccnj, Blnk, Dntt, Tm9sf3, Pik3ap1, Lcor, Slit1, Arhgap19, Rrp12, Pgam1, Exosc1, Mms19, Ubtd1, Morn4, Pi4k2a, Zfyve27, Golga7b, Crtac1, R3hcc1l, Loxl4, Pyroxd2, Hps1, Hpse2, Dnmbp, Cyp2c23, Erlin1, Cwf19l1, Bloc1s2, Scd3, Scd4, Scd1, Wnt8b, Sec31b, Ndufb8, Hif1an, Slf2, Sema4g, Mrpl43, Lzts2, Kazald1, Btrc, Fbxw4, Kcnip2, Pprc1, Elovl3, Gbf1, Psd, Sufu, Trim8, Sfxn2, Wbp1l, Cyp17a1, Borcs7, As3mt, Cnnm2, Nt5c2, Ina, Pcgf6, Pdcd11, Neurl1a, Sh3pxd2a, Stn1, Slk, Col17a1, Gsto1, Cfap58, Sorcs3, Sorcs1, Xpnpep1, Mxi1, Rbm20, Pdcd4, Shoc2, Tectb, Acsl5, Vti1a, Tcf7l2, Nrap, Plekhs1, Ablim1, Fam160b1, Trub1, Atrnl1, Gfra1, Pnliprp1, Hspa12a, Eno4, Shtn1, Slc18a2, Pdzd8, Rab11fip2, Cacul1, Fam45a, Sfxn4, Grk5, Csf2ra* | *Loxl4, Pyroxd2, Hps1, Lzts2, Trim8* |
| *Qlh14* | **Lineage⁻ (Lin⁻)** | Chr. 12: 106.34 – 107.94Mb | WSB – low | *Rit1* | |

Supplemental Table 4: Relationships between mapped QTL and all cellular phenotypes measured in the lungs of CC-F1 animals. Bold – $p < 0.05$, per test; Bold + Italic – FDR $p < 0.05$

| PHENOTYPES | *Qlh1* | *Qlh2* | *Qlh3* | *Qlh4* | *Qlh5* | *Qlh6* | *Qlh7* | *Qlh8* | *Qlh9* | *Qlh10* | *Qlh11* | *Qlh12* | *Qlh13* | *Qlh14* | *Qlh15* | *Qlh16* |
|---|---|---|---|---|---|---|---|---|---|---|---|---|---|---|---|---|
| CD103+, CD205+ DCs | 0.3277 | 0.4924 | 0.4879 | ***6.36E-06*** | 0.5259 | 0.9460 | 0.6732 | 0.9921 | 0.3674 | 0.7278 | 0.7766 | 0.5069 | 0.4617 | 0.5090 | 0.3685 | 0.5908 |
| CD326+ DCs | 0.3857 | 0.8703 | 0.6396 | **0.0361** | 0.5828 | **0.0029** | 0.2202 | 0.7710 | 0.9915 | 0.6946 | 0.8060 | 0.1826 | 0.3883 | 0.1514 | 0.2037 | 0.8998 |
| CD326+ DCs (prop. of Lin-) | 0.2505 | 0.8363 | 0.8837 | 0.5219 | 0.7223 | **0.0077** | 0.0814 | 0.3645 | 0.9484 | 0.2510 | 0.4835 | 0.7486 | 0.4929 | 0.5649 | 0.8828 | 0.7634 |
| CD103+, CD205+ DCs (prop. of Lin-) | 0.2309 | 0.8029 | 0.4899 | ***7.05E-04*** | 0.4540 | 0.8739 | 0.6154 | 0.9615 | 0.3457 | 0.9678 | 0.7799 | 0.4468 | 0.3390 | 0.6820 | 0.3469 | 0.6787 |
| Ly6C- monos/macs (prop. of Ly6C$^{+/lo/-}$) | **0.0294** | 0.1162 | 0.1337 | 0.2924 | 0.6389 | **0.0213** | ***1.62E-04*** | **2.56E-07** | 0.6336 | **0.0053** | ***2.05E-07*** | 0.7616 | 0.4200 | 0.4408 | **0.0263** | ***0.0012*** |
| Eosinophils | 0.6178 | ***0.0015*** | 0.2523 | 0.6046 | 0.5713 | 0.5116 | 0.4954 | 0.1736 | 0.7792 | 0.1346 | 0.1496 | ***5.58E-06*** | 0.6585 | **0.0022** | 0.7549 | 0.3921 |
| CD103+ DCs | **0.0508** | 0.4648 | 0.5220 | 0.0586 | 0.4966 | 0.4969 | 0.3308 | 0.8593 | 0.4518 | 0.8074 | 0.5566 | 0.3287 | 0.4235 | 0.8608 | 0.4000 | 0.7092 |
| NK cells | 0.7494 | 0.8602 | 0.3080 | 0.0708 | 0.8076 | 0.1803 | **0.0310** | 0.4055 | 0.5751 | 0.1548 | 0.3394 | 0.1074 | 0.9812 | 0.5885 | 0.3149 | 0.2147 |
| Lineage Negative | 0.6459 | 0.5132 | 0.1516 | 0.1197 | **0.0161** | 0.9040 | 0.6674 | 0.3249 | 0.7910 | **0.0130** | 0.0658 | **0.0060** | 0.5407 | **0.0010** | **8.66E-04** | 0.4868 |
| Lineage Positive | 0.5526 | 0.8070 | 0.1263 | 0.1157 | **0.0170** | 0.9209 | 0.5154 | 0.3230 | 0.7471 | **0.0121** | 0.0956 | **0.0045** | 0.5411 | ***0.0015*** | ***6.45E-04*** | 0.5168 |
| pDCs | 0.9485 | 0.5669 | 0.9780 | **0.0367** | 0.6667 | 0.3920 | 0.1813 | **3.25E-07** | 0.1257 | **1.17E-04** | ***7.94E-04*** | 0.1488 | 0.2102 | 0.6666 | 0.3897 | 0.0576 |
| Ly6C+ monos/macs (prop. of Ly6C$^{+/lo/-}$) | 0.1243 | 0.1112 | 0.1371 | **0.0393** | 0.7063 | 0.2113 | ***0.0010*** | **1.57E-10** | 0.7097 | ***1.31E-04*** | ***2.61E-08*** | 0.8749 | 0.6102 | 0.9904 | 0.1155 | 0.3438 |
| CD8+ DCs | **0.0290** | 0.9001 | 0.3418 | 0.7620 | **0.0136** | 0.4857 | 0.9029 | 0.3010 | 0.7773 | **0.0202** | 0.2299 | 0.5467 | 0.4882 | 0.6624 | 0.2054 | ***7.76E-04*** |
| NKT cells (prop. of CD3+ T cells) | 0.1831 | 0.8444 | 0.1269 | 0.0638 | 0.2268 | 0.5192 | 0.4648 | 0.5392 | 0.4646 | 0.9322 | 0.9430 | 0.5522 | 0.3328 | 0.1320 | 0.1324 | **0.0443** |
| Ly6C$^{-}$ monos/macs | **0.0025** | 0.0950 | 0.1036 | 0.7209 | 0.8984 | **0.0189** | 0.2586 | **0.0076** | 0.3344 | 0.0574 | **1.03E-06** | 0.3386 | 0.7426 | 0.5193 | 0.6370 | **0.0439** |
| CD11b+ cells | 0.1984 | 0.5286 | 0.0943 | 0.0969 | 0.8933 | 0.5605 | 0.2718 | 0.9997 | 0.7670 | 0.8262 | 0.7750 | 0.1044 | 0.2573 | **0.0074** | **0.0038** | 0.7522 |
| CD11b+ cells (prop. of Lin-) | 0.1984 | 0.5286 | 0.0943 | 0.0969 | 0.8933 | 0.5605 | 0.2718 | 0.9997 | 0.7670 | 0.8262 | 0.7750 | 0.1044 | 0.2573 | **0.0074** | **0.0038** | 0.7522 |
| Ly6C$^{lo}$ monos/macs (prop. of Ly6C$^{+/lo/-}$) | 0.0960 | 0.5000 | 0.5492 | 0.9180 | 0.7752 | **0.0456** | **0.0069** | 0.3121 | 0.7583 | 0.3820 | **0.0194** | 0.8159 | 0.6835 | 0.3015 | 0.1656 | **1.91E-04** |
| Neutrophils | 0.6555 | 0.5544 | 0.5845 | 0.9476 | 0.4529 | 0.5397 | 0.7976 | 0.8433 | 0.4442 | 0.3663 | 0.4627 | 0.1175 | **0.0073** | 0.4460 | 0.6560 | 0.4089 |
| CD8+ Ts | 0.1926 | ***0.0011*** | **0.0107** | **0.0098** | **2.92E-05** | 0.5242 | 0.5139 | 0.6926 | 0.4410 | **0.0111** | 0.0997 | 0.2311 | 0.1360 | 0.8579 | **0.0130** | 0.1693 |
| Inflammatory DCs | **0.0193** | 0.1305 | 0.4075 | 0.5631 | 0.0949 | 0.7177 | 0.8350 | 0.8447 | **0.0374** | 0.1287 | 0.0611 | ***2.64E-04*** | 0.0845 | **0.0212** | 0.9059 | 0.9900 |
| CD3+ T cells | **0.0114** | 0.6881 | **0.0039** | **0.0001** | 0.0882 | 0.0985 | 0.3807 | 0.4665 | 0.3590 | 0.8012 | 0.2098 | 0.5712 | **0.0075** | 0.9426 | **0.0046** | 0.2265 |
| CD103+ DCs (prop. of Lin-) | **0.0144** | 0.8143 | 0.6191 | 0.2560 | 0.3518 | 0.5134 | 0.2616 | 0.9685 | 0.4322 | 0.8923 | 0.8422 | 0.3273 | 0.3037 | 0.9800 | 0.5227 | 0.7137 |
| CD11b+ DCs (prop. of Lin-) | 0.2995 | 0.8600 | 0.2695 | 0.0876 | 0.2614 | 0.7436 | 0.4844 | 0.9880 | 0.4510 | 0.6965 | 0.1884 | 0.4573 | 0.5500 | 0.2384 | 0.0676 | 0.4470 |
| B cells | 0.1259 | 0.1885 | 0.2000 | 0.6446 | 0.9284 | 0.3845 | 0.3655 | 0.6727 | 0.6001 | 0.6820 | 0.6790 | 0.3817 | **0.0426** | **0.0245** | 0.2382 | 0.2404 |
| CD8+ T cells (prop. of CD3+ T cells) | 0.4482 | **1.13E-04** | **0.0171** | 0.5035 | ***1.71E-05*** | 0.8549 | 0.8189 | 0.8190 | 0.0514 | **0.0074** | **0.0109** | 0.1952 | 0.0741 | 0.8455 | **0.0298** | 0.3713 |
| CD205+ DCs (prop. of Lin-) | 0.6275 | **0.0061** | 0.7746 | 0.7208 | 0.1729 | **0.0501** | 0.9013 | 0.6284 | 0.4256 | 0.5066 | 0.3502 | 0.1178 | 0.5351 | 0.8759 | 0.9038 | 0.9916 |
| CD103-, CD205- DCs (prop. of Lin-) | 0.0861 | 0.2491 | 0.8384 | 0.6876 | 0.5908 | 0.2856 | ***1.24E-08*** | 0.5059 | 0.9049 | 0.4750 | 0.6004 | 0.9454 | 0.6486 | 0.1443 | 0.3051 | 0.6493 |
| CD11c+ cells | 0.1600 | 0.7446 | 0.1731 | 0.1018 | 0.2837 | 0.2068 | **0.0408** | 0.9760 | 0.5009 | 0.3056 | 0.7022 | **0.0098** | 0.5744 | 0.1068 | **0.0355** | 0.6719 |
| Gr1$^{lo}$ monos/macs (prop. of Gr1$^{+/lo/-}$) | 0.7544 | 0.7460 | 0.1139 | 0.1942 | 0.7699 | 0.8896 | 0.1211 | 0.6483 | 0.3278 | 0.5408 | 0.5233 | ***6.03E-04*** | 0.1061 | ***3.66E-07*** | ***4.56E-07*** | 0.0859 |
| MHCII+ DCs (prop. of CD11b+) | 0.4917 | 0.5896 | 0.1299 | 0.2949 | 0.8451 | 0.5865 | 0.7807 | 0.2033 | 0.7734 | 0.7360 | **0.0148** | 0.2120 | 0.0518 | 0.2787 | 0.9658 | 0.7544 |
| Ly6C$^{+}$ monos/macs | 0.1059 | 0.1109 | 0.4425 | 0.1039 | 0.8575 | 0.2416 | **1.19E-18** | **2.25E-08** | 0.9698 | **0.0034** | **0.0110** | 0.9011 | 0.9242 | 0.7076 | 0.2018 | 0.0873 |
| NK T cells | 0.2357 | 0.8800 | 0.3569 | 0.5322 | 0.8871 | 0.6806 | 0.5171 | 0.8333 | 0.3823 | 0.6651 | 0.6076 | 0.4997 | 0.3183 | 0.4150 | 0.3359 | **0.0328** |
| Total Leukocytes (CD45+) | 0.0911 | 0.4600 | 0.7242 | 0.8160 | 0.8465 | 0.6835 | ***8.91E-05*** | 0.5163 | 0.9506 | 0.8611 | 0.2336 | **0.0120** | 0.8795 | 0.0665 | **0.0418** | 0.5978 |
| CD11b+ DCM | 0.5497 | 0.1025 | 0.2718 | ***2.67E-07*** | **0.0370** | **0.0262** | 0.5665 | 0.0598 | 0.5825 | 0.1360 | 0.4137 | 0.4278 | 0.7474 | 0.6641 | 0.1731 | **0.0304** |
| Gr1$^{lo}$ monos/macs | 0.6900 | 0.9078 | 0.7023 | 0.9536 | 0.7923 | 0.6023 | 0.5926 | 0.8277 | 0.6324 | 0.5105 | 0.4698 | ***2.91E-05*** | 0.1571 | ***2.23E-25*** | ***5.64E-04*** | **0.0091** |

| | | | | | | | | | | | | | | | | |
|---|---|---|---|---|---|---|---|---|---|---|---|---|---|---|---|---|
| CD4+ T cells | **7.56E-04** | 0.4850 | **0.0092** | 0.1584 | 0.2324 | 0.1438 | 0.2740 | 0.6347 | 0.1831 | 0.2146 | **0.0038** | 0.7974 | ***2.62E-04*** | 0.3906 | **0.0122** | 0.5839 |
| MHCII+ DCs | **0.0457** | 0.7231 | 0.2358 | ***1.82E-05*** | **0.0323** | 0.0872 | 0.8459 | 0.1136 | 0.9497 | **0.0135** | **0.0090** | 0.5084 | **0.0167** | 0.6159 | 0.4261 | 0.5331 |
| CD205+ DCs | 0.5263 | 0.6583 | 0.4592 | 0.6130 | 0.4051 | 0.0679 | 0.8732 | 0.9357 | 0.5797 | 0.6391 | 0.3215 | 0.0552 | 0.7677 | 0.6066 | 0.2799 | 0.8276 |
| CD11b- cells (prop. of CD11b$^{+/-}$) | 0.4152 | 0.1033 | 0.4043 | 0.2518 | 0.4910 | 0.3770 | 0.5101 | 0.4692 | 0.6974 | 0.3795 | 0.2476 | 0.1659 | 0.2749 | 0.5018 | 0.1409 | 0.7536 |
| CD11b- cells | 0.2052 | 0.7501 | 0.3558 | 0.1916 | 0.3210 | 0.2762 | **0.0354** | 0.9743 | 0.4283 | 0.3656 | 0.5974 | **0.0302** | 0.7469 | 0.4192 | 0.1696 | 0.5186 |
| CD4-, CD8- T cells (prop. of CD3+ T cells) | ***6.78E-04*** | 0.4690 | **0.0418** | 0.7275 | 0.9419 | 0.3294 | **0.0150** | 0.2862 | 0.5389 | 0.1676 | 0.2169 | 0.6343 | 0.6010 | 0.1014 | 0.0654 | 0.8473 |
| CD4+ -to- CD8+ T cell ratio | **0.0221** | 0.1026 | 0.3748 | 0.6149 | 0.0569 | 0.3693 | **0.0120** | 0.2337 | 0.2242 | **0.0079** | **0.0024** | 0.1980 | 0.1541 | 0.1806 | 0.3897 | 0.8585 |
| CD8+ -to- CD4+ T cell ratio | **0.0221** | 0.1026 | 0.3748 | 0.6149 | 0.0569 | 0.3693 | **0.0120** | 0.2337 | 0.2242 | **0.0079** | **0.0024** | 0.1980 | 0.1541 | 0.1806 | 0.3897 | 0.8585 |
| CD11b+ DCs | 0.0876 | 0.6772 | 0.1184 | 0.2171 | 0.7722 | 0.6446 | 0.3653 | 0.9637 | 0.6467 | 0.9426 | 0.6420 | 0.1645 | 0.3778 | **0.0157** | **0.0069** | 0.7336 |
| CD103-, CD205- DCs | 0.1184 | 0.9020 | 0.7719 | 0.4507 | 0.8448 | 0.2731 | **1.89E-12** | 0.4948 | 0.8298 | 0.1097 | 0.9977 | 0.1995 | 0.7536 | 0.5216 | 0.5023 | 0.3029 |
| CD11b+ cells (prop. of Lin-) | 0.1810 | **0.0100** | 0.6782 | 0.3114 | 0.0869 | 0.1626 | 0.0794 | 0.7249 | 0.3092 | 0.4899 | 0.4266 | 0.1059 | 0.4956 | 0.7764 | 0.8932 | 0.9421 |
| Ly6C$^{lo}$ monos/macs | 0.6695 | 0.6927 | 0.9282 | 0.1888 | 0.2352 | 0.4858 | **3.45E-04** | 0.1911 | 0.7783 | 0.3436 | 0.5698 | 0.7893 | 0.7565 | 0.6483 | 0.5203 | ***7.33E-05*** |
| Gr1$^{-}$ monos/macs | 0.5673 | 0.9915 | 0.1935 | 0.1335 | **0.0051** | 0.6165 | 0.8583 | 0.2915 | 0.5688 | 0.1790 | 0.0728 | 0.6241 | **0.0044** | 0.5034 | 0.0845 | 0.1678 |
| CD4+ T cells (prop. of CD3+ T cells) | **0.0125** | 0.2561 | 0.4932 | 0.7286 | 0.1759 | 0.3517 | **0.0175** | 0.2175 | 0.1760 | **0.0110** | **0.0071** | 0.3226 | 0.1550 | 0.0934 | 0.5168 | 0.9424 |
| CD4$^{-}$, CD8$^{-}$ T cells | **0.0310** | 0.5735 | 0.1306 | 0.6437 | 0.9414 | 0.6417 | **0.0252** | 0.3517 | 0.7608 | 0.2049 | 0.4681 | 0.6213 | 0.6119 | 0.1206 | 0.1998 | 0.8717 |
| CD11c$^{+}$ cells (prop. of Lin$^{-}$) | 0.2310 | **0.0172** | 0.3735 | 0.2562 | **0.0247** | 0.0851 | 0.0564 | 0.6074 | 0.4329 | 0.3312 | 0.2704 | 0.0576 | 0.3040 | 0.6938 | 0.4364 | 0.9838 |
| Alveolar macrophages | 0.1713 | 0.9689 | 0.0757 | ***8.20E-05*** | 0.4838 | **1.30E-04** | 0.1403 | 0.2244 | 0.5964 | 0.4895 | 0.7187 | 0.4676 | 0.7150 | 0.8215 | **0.0440** | 0.4911 |
| CD8+ DCs (prop. of Lin$^{-}$) | 0.1577 | 0.8636 | 0.5490 | 0.4158 | 0.8996 | 0.4286 | 0.7853 | 0.9380 | 0.6842 | 0.9681 | 0.2648 | ***7.66E-05*** | ***2.13E-06*** | **0.0076** | ***0.0017*** | 0.0952 |

| PHENOTYPES | *Qlh17* | *Qlh18* | *Qlh19* | *Qlh20* | *Qlh21* | *Qlh22* | *Qlh23* | *Qlh24* | *Qlh25* | *Qlh26* | *Qlh27* | *Qlh28* |
|---|---|---|---|---|---|---|---|---|---|---|---|---|
| CD103+, CD205+ DCs | 0.5069 | 0.6420 | 0.8271 | **0.0170** | 0.2677 | 0.9005 | 0.1324 | 0.8147 | 0.4084 | 0.9849 | 0.4385 | ***0.0013*** |
| CD326+ DCs | 0.1826 | 0.7343 | 0.5306 | 0.3253 | 0.1020 | 0.6163 | 0.8100 | 0.8979 | 0.3275 | 0.7430 | 0.4607 | **0.0032** |
| CD326+ DCs (prop. of Lin-) | 0.7486 | 0.3142 | 0.8461 | 0.3012 | 0.1106 | 0.9713 | 0.8357 | 0.9183 | 0.5468 | 0.7813 | 0.4194 | **0.0050** |
| CD103+, CD205+ DCs (prop. of Lin-) | 0.4468 | 0.2939 | 0.8831 | **0.0026** | 0.5037 | 0.8556 | 0.1678 | 0.8066 | 0.3249 | 0.9765 | 0.5148 | **0.0105** |
| Ly6C- monos/macs (prop. of Ly6C$^{+/lo/-}$) | 0.7616 | 0.2248 | 0.4894 | 0.5445 | 0.9593 | 0.1064 | 0.2654 | 0.4242 | 0.3526 | **0.0204** | 0.4939 | 0.0859 |
| Eosinophils | ***5.58E-06*** | 0.6173 | **0.0098** | 0.5670 | 0.6006 | 0.7384 | ***7.59E-05*** | 0.2500 | 0.5272 | 0.7105 | 0.1210 | 0.1246 |
| CD103+ DCs | 0.3287 | 0.4797 | 0.6274 | 0.6621 | 0.7112 | 0.4275 | 0.1289 | 0.1812 | 0.6834 | 0.9197 | 0.4291 | 0.1410 |
| NK cells | 0.1074 | 0.7440 | **0.0274** | **0.0393** | 0.6421 | 0.2242 | 0.6476 | 0.7043 | **0.0227** | 0.9077 | 0.5300 | 0.1453 |
| Lineage Negative | **0.0060** | 0.1536 | 0.1754 | 0.6995 | 0.5790 | 0.8364 | 0.6171 | 0.6579 | 0.4356 | 0.2576 | 0.8481 | 0.1615 |
| Lineage Positive | **0.0045** | 0.1371 | 0.3473 | 0.6238 | 0.5335 | 0.9396 | 0.4611 | 0.6735 | 0.3923 | 0.2928 | 0.8082 | 0.1746 |
| pDCs | 0.1488 | 0.3808 | 0.8259 | 0.0626 | 0.1632 | 0.1415 | 0.4481 | 0.9276 | 0.1589 | **0.0488** | 0.8716 | 0.1869 |
| Ly6C+ monos/macs (prop. of Ly6C$^{+/lo/-}$) | 0.8749 | 0.5881 | 0.1892 | 0.2403 | 0.2648 | 0.1021 | 0.2740 | 0.2487 | **0.0155** | **0.0343** | 0.6344 | 0.1952 |
| CD8+ DCs | 0.5467 | 0.8431 | 0.7641 | 0.3451 | 0.8577 | 0.3292 | 0.6015 | 0.6017 | 0.5140 | 0.7273 | **0.0030** | 0.1997 |
| NKT cells (prop. of CD3+ T cells) | 0.5522 | 0.8694 | 0.2247 | 0.2511 | 0.6989 | 0.4624 | 0.7923 | 0.9536 | 0.7564 | 0.6292 | 0.1032 | 0.2350 |
| Ly6C$^{-}$ monos/macs | 0.3386 | 0.1445 | 0.1822 | 0.4044 | 0.4134 | ***0.0016*** | **0.0419** | 0.3076 | 0.5163 | **0.0561** | 0.4768 | 0.2475 |
| CD11b+ cells | 0.1044 | 0.1662 | **0.0151** | 0.9077 | 0.3236 | 0.7576 | 0.2823 | 0.9300 | **0.0305** | 0.1499 | 0.4373 | 0.2572 |
| CD11b+ cells (prop. of Lin-) | 0.1044 | 0.1662 | **0.0151** | 0.9077 | 0.3236 | 0.7576 | 0.2823 | 0.9300 | **0.0305** | 0.1499 | 0.4373 | 0.2572 |
| Ly6C$^{lo}$ monos/macs (prop. of Ly6C$^{+/lo/-}$) | 0.8159 | 0.5635 | 0.0975 | 0.9042 | 0.3054 | 0.5997 | 0.6357 | 0.6967 | 0.9395 | 0.2167 | **0.0448** | 0.2917 |
| Neutrophils | 0.1175 | 0.4101 | 0.5746 | 0.2289 | 0.3102 | 0.3903 | 0.4833 | **0.0100** | 0.7317 | 0.9814 | **0.0134** | 0.3221 |
| CD8+ Ts | 0.2311 | 0.5299 | 0.9725 | 0.4852 | 0.4863 | 0.8175 | 0.7676 | 0.9272 | 0.4097 | 0.1079 | 0.1181 | 0.3588 |
| Inflammatory DCs | ***2.64E-04*** | 0.3222 | 0.1833 | 0.6877 | 0.1890 | 0.1855 | **3.86E-04** | **0.0791** | **0.0491** | **0.0855** | 0.3039 | 0.3675 |
| CD3+ T cells | 0.5712 | 0.9496 | 0.9319 | 0.4231 | 0.2510 | 0.4120 | 0.4514 | 0.4035 | 0.1117 | 0.5569 | 0.1274 | 0.3688 |

| | | | | | | | | | | | | |
|---|---|---|---|---|---|---|---|---|---|---|---|---|
| CD103+ DCs (prop. of Lin-) | 0.3273 | 0.4928 | 0.5388 | 0.3683 | 0.9623 | 0.6843 | 0.0902 | 0.1466 | 0.6366 | 0.9424 | 0.4650 | 0.3901 |
| CD11b+ DCs (prop. of Lin-) | 0.4573 | 0.1373 | **0.0127** | 0.2185 | 0.4415 | 0.6654 | **0.0093** | 0.7077 | 0.1268 | 0.7472 | 0.4161 | 0.3940 |
| B cells | 0.3817 | 0.3470 | 0.5239 | 0.1526 | **0.0461** | 0.4877 | 0.2119 | **0.0630** | 0.5783 | 0.9596 | 0.3978 | 0.4544 |
| CD8+ T cells (prop. of CD3+ T cells) | 0.1952 | 0.2087 | 0.9850 | 0.7359 | 0.9064 | 0.9774 | 0.4819 | 0.7334 | 0.2646 | **0.0186** | 0.5541 | 0.4826 |
| CD205+ DCs (prop. of Lin-) | 0.1178 | 0.1922 | 0.7828 | 0.4706 | 0.7770 | 0.4583 | 0.3534 | 0.6245 | 0.4813 | 0.3562 | 0.9735 | 0.4893 |
| CD103-, CD205- DCs (prop. of Lin-) | 0.9454 | 0.8215 | ***2.13E-07*** | 0.6881 | 0.4327 | 0.6274 | 0.5265 | 0.3573 | 0.8271 | 0.7595 | 0.9801 | 0.4978 |
| CD11c+ cells | **0.0098** | **0.0313** | 0.2287 | 0.4943 | 0.8056 | 0.7822 | 0.2305 | 0.6813 | 0.6867 | 0.8534 | 0.9004 | 0.5069 |
| Gr1$^{lo}$ monos/macs (prop. of Gr1$^{+/lo/-}$) | ***6.03E-04*** | 0.5422 | ***5.50E-07*** | 0.3944 | ***1.70E-04*** | 0.9962 | 0.3536 | 0.7204 | **0.0462** | 0.6676 | 0.9886 | 0.5083 |
| MHCII+ DCs (prop. of CD11b+) | 0.2120 | 0.2120 | 0.5264 | 0.2408 | 0.5574 | 0.3329 | 0.1613 | 0.1802 | ***4.04E-04*** | **0.0337** | 0.7868 | 0.5311 |
| Ly6C$^{+}$ monos/macs | 0.9011 | 0.4421 | ***6.57E-05*** | 0.7224 | **0.0066** | 0.6112 | 0.9484 | 0.3316 | 0.5888 | 0.6775 | 0.9918 | 0.5326 |
| NK T cells | 0.4997 | 0.7672 | 0.1429 | 0.3580 | 0.4878 | 0.8392 | 0.6769 | 0.4777 | 0.2881 | 0.6353 | 0.1269 | 0.5368 |
| Total Leukocytes (CD45+) | **0.0120** | 0.2953 | 0.0520 | 0.2598 | 0.4696 | 0.5378 | 0.5342 | 0.1204 | 0.8680 | 0.8242 | 0.5382 | 0.5622 |
| CD11b+ DCM | 0.4278 | 0.7157 | 0.3720 | 0.9447 | 0.8413 | 0.0880 | **0.0050** | **0.0314** | ***5.25E-05*** | **0.0285** | 0.5003 | 0.5660 |
| Gr1$^{lo}$ monos/macs | ***2.91E-05*** | 0.3945 | ***2.51E-10*** | 0.1767 | **0.0028** | 0.4073 | 0.3006 | 0.7325 | 0.5761 | 0.1263 | 0.6836 | 0.5842 |
| CD4+ T cells | 0.7974 | 0.7039 | 0.1907 | 0.0564 | 0.1975 | 0.1688 | 0.5776 | 0.9147 | 0.1651 | **0.0390** | 0.3671 | 0.6006 |
| MHCII+ DCs | 0.5084 | 0.4224 | 0.6868 | 0.6844 | 0.3753 | 0.6258 | ***5.09E-04*** | 0.1338 | **4.66E-04** | **1.14E-04** | 0.6637 | 0.6936 |
| CD205+ DCs | 0.0552 | 0.0512 | 0.8702 | 0.3029 | 0.8886 | 0.8522 | 0.3319 | 0.9832 | 0.7098 | 0.6825 | 0.9546 | 0.6950 |
| CD11b- cells (prop. of CD11b$^{+/-}$) | 0.1659 | 0.0839 | 0.2665 | 0.9169 | 0.6873 | 0.4531 | ***0.0015*** | 0.5616 | **0.0295** | 0.2560 | 0.4547 | 0.7076 |
| CD11b- cells | **0.0302** | **0.0311** | 0.3374 | 0.5164 | 0.9906 | 0.8469 | 0.0834 | 0.7299 | 0.7121 | 0.6927 | 0.8846 | 0.7089 |
| CD4-, CD8- T cells (prop. of CD3+ T cells) | 0.6343 | 0.8435 | **0.0309** | ***0.0019*** | 0.1453 | 0.3444 | 0.7828 | 0.3559 | 0.8088 | 0.3267 | 0.7631 | 0.7307 |
| CD4+ -to- CD8+ T cell ratio | 0.1980 | 0.5805 | **0.0425** | **0.0203** | 0.1422 | 0.2729 | 0.6445 | 0.7617 | 0.5281 | **0.0053** | 0.7402 | 0.7320 |
| CD8+ -to- CD4+ T cell ratio | 0.1980 | 0.5805 | **0.0425** | **0.0203** | 0.1422 | 0.2729 | 0.6445 | 0.7617 | 0.5281 | **0.0053** | 0.7402 | 0.7320 |
| DblNeg | 0.1645 | 0.0568 | **5.38E-04** | 0.1896 | 0.3462 | 0.3738 | 0.0641 | 0.6859 | **0.0723** | 0.2618 | 0.3812 | 0.7754 |
| CD103-, CD205- DCs | 0.1995 | 0.2671 | ***2.69E-05*** | 0.5171 | 0.4689 | 0.5376 | 0.4993 | 0.6441 | 0.8711 | 0.8477 | 0.9441 | 0.7932 |
| CD11b+ cells (prop. of Lin-) | 0.1059 | 0.1904 | 0.1477 | 0.8735 | 0.7673 | 0.6517 | 0.0554 | 0.2714 | 0.5847 | 0.3705 | 0.9295 | 0.8353 |
| Ly6C$^{lo}$ monos/macs | 0.7893 | 0.7298 | **0.0146** | 0.1576 | 0.8116 | 0.0950 | 0.6574 | 0.7802 | 0.5464 | 0.4559 | 0.2283 | 0.8468 |
| Gr1$^{-}$ monos/macs | 0.6241 | 0.8120 | 0.1833 | 0.1446 | 0.2971 | 0.1794 | 0.2755 | 0.2730 | 0.1058 | 0.1198 | 0.2878 | 0.9159 |
| CD4+ T cells (prop. of CD3+ T cells) | 0.3226 | 0.5975 | **0.0484** | **0.0065** | 0.1572 | 0.4090 | 0.7482 | 0.6312 | 0.4843 | **0.0139** | 0.6643 | 0.9291 |
| CD4$^{-}$, CD8$^{-}$ T cells | 0.6213 | 0.7366 | 0.0581 | **0.0213** | 0.3602 | 0.3370 | 0.3971 | 0.1838 | 0.7314 | 0.2977 | 0.9091 | 0.9586 |
| CD11c$^{+}$ cells (prop. of Lin$^{-}$) | 0.0576 | 0.1867 | 0.1900 | 0.8403 | 0.5337 | 0.6444 | 0.2651 | 0.1908 | 0.7202 | 0.3555 | 0.8554 | 0.9605 |
| Alveolar macrophages | 0.4676 | 0.8369 | 0.4658 | 0.3642 | 0.4303 | 0.1615 | 0.1491 | 0.5899 | 0.3171 | 0.7349 | 0.0823 | 0.9629 |
| CD8+ DCs (prop. of Lin$^{-}$) | ***7.66E-05*** | 0.3422 | 0.0824 | 0.2816 | **0.0479** | 0.3543 | 0.7256 | 0.6178 | 0.7606 | 0.5691 | ***9.26E-05*** | 0.9909 |

Supplemental Table 5: Relationships between mapped QTL and influenza A virus induced disease phenotypes. Bold – $p < 0.05$, per test

| IAV PHENOTYPE | *Qlh1* | *Qlh2* | *Qlh3* | *Qlh4* | *Qlh5* | *Qlh6* | *Qlh7* | *Qlh8* | *Qlh9* | *Qlh10* | *Qlh11* | *Qlh12* | *Qlh13* | *Qlh14* |
|---|---|---|---|---|---|---|---|---|---|---|---|---|---|---|
| % starting weight - D1 | ***0.0926*** | 0.9498 | 0.7578 | 0.8477 | 0.2643 | 0.9221 | 0.2164 | ***0.1853*** | 0.4624 | 0.2917 | 0.9857 | 0.4178 | 0.3295 | 0.5553 |
| % starting weight - D2 | ***1.66E-11*** | ***0.0874*** | 0.7040 | 0.5614 | 0.7487 | ***0.1222*** | 0.4749 | 0.9454 | 0.5926 | 0.9342 | 0.9381 | 0.9319 | 0.7817 | 0.8926 |
| % starting weight - D3 | ***2.61E-07*** | ***0.0330*** | 0.8236 | 0.6870 | 0.8062 | 0.3111 | 0.5836 | 0.9275 | 0.5849 | 0.9266 | 0.8408 | 0.5288 | 0.6178 | 0.8709 |
| % starting weight - D4 | ***1.42E-06*** | ***0.0141*** | 0.8580 | 0.3693 | 0.8440 | 0.3125 | ***0.0047*** | 0.4580 | 0.3464 | 0.5047 | 0.5134 | 0.5462 | ***0.1777*** | 0.7552 |
| % starting weight - D5 | ***0.0055*** | ***0.0222*** | 0.3295 | 0.4378 | 0.8826 | 0.6915 | ***0.0442*** | 0.2559 | 0.3017 | 0.3635 | 0.3276 | 0.4686 | ***0.1368*** | 0.7482 |
| % starting weight - D6 | 0.3209 | ***0.1045*** | 0.9415 | 0.6594 | 0.9788 | 0.7047 | 0.9471 | 0.9496 | 0.9141 | 0.9869 | 0.9972 | 0.5841 | 0.6516 | 0.9808 |
| % starting weight - D7 | ***0.0340*** | ***0.0371*** | ***0.1893*** | 0.7614 | 0.9189 | 0.8280 | ***0.1022*** | 0.2606 | ***0.0507*** | 0.6927 | ***0.1392*** | 0.4155 | 0.4272 | 0.8601 |
| % starting weight - D8 | ***0.0028*** | ***0.0125*** | 0.4569 | 0.6872 | 0.7100 | 0.4634 | ***0.1053*** | 0.3835 | ***0.0406*** | 0.8542 | 0.2366 | ***0.1179*** | 0.4854 | 0.5223 |
| % starting weight - D9 | ***0.0120*** | ***0.0064*** | ***0.1708*** | 0.4409 | 0.8243 | 0.3559 | ***0.1147*** | 0.3398 | ***0.0491*** | 0.6050 | ***0.1896*** | ***0.1076*** | 0.5922 | 0.3436 |
| % starting weight - D10 | 0.6082 | 0.6968 | ***0.0008*** | ***0.0010*** | 0.5534 | 0.3691 | ***0.0244*** | 0.9441 | 0.8212 | ***0.0461*** | ***0.1110*** | 0.2917 | 0.5305 | 0.9660 |
| Day of lowest weight | ***0.1880*** | ***0.0380*** | ***0.1794*** | 0.3484 | 0.8152 | 0.3515 | 0.3135 | 0.3968 | ***0.1076*** | 0.5783 | 0.4333 | ***0.1426*** | 0.4443 | 0.5500 |
| Lowest weight reached | 0.2604 | 0.2927 | 0.3496 | 0.5749 | 0.9082 | ***0.0046*** | 0.4821 | 0.9673 | 0.4422 | 0.9693 | 0.7813 | 0.2069 | 0.7299 | 0.2182 |

| IAV PHENOTYPE | *Qlh15* | *Qlh16* | *Qlh17* | *Qlh18* | *Qlh19* | *Qlh20* | *Qlh21* | *Qlh22* | *Qlh23* | *Qlh24* | *Qlh25* | *Qlh26* | *Qlh27* | *Qlh28* |
|---|---|---|---|---|---|---|---|---|---|---|---|---|---|---|
| % starting weight - D1 | 0.4988 | 0.3889 | 0.4178 | 0.2672 | 0.4952 | 0.3306 | 0.3472 | 0.6019 | 0.4138 | ***0.1649*** | 0.4496 | 0.4799 | 0.6002 | 0.6346 |
| % starting weight - D2 | 0.6188 | 0.7451 | 0.9319 | 0.2520 | ***0.0711*** | ***0.1404*** | 0.9059 | ***0.1165*** | 0.4691 | 0.4524 | 0.8149 | 0.5550 | 0.7128 | ***0.0973*** |
| % starting weight - D3 | 0.8934 | 0.9272 | 0.5288 | 0.4352 | ***0.1119*** | 0.5100 | 0.9824 | 0.3635 | 0.4763 | 0.6414 | 0.8589 | 0.3955 | 0.9760 | 0.2312 |
| % starting weight - D4 | 0.6847 | 0.7727 | 0.5462 | ***0.1201*** | ***0.0415*** | 0.5575 | 0.4827 | ***0.0418*** | 0.5993 | 0.9218 | 0.6263 | 0.4374 | 0.8181 | 0.2610 |
| % starting weight - D5 | 0.8481 | 0.4055 | 0.4686 | ***0.0142*** | ***0.1282*** | 0.2685 | 0.7586 | ***0.0136*** | 0.4807 | 0.6868 | ***0.1782*** | 0.5507 | 0.9828 | 0.5624 |
| % starting weight - D6 | 0.9132 | 0.2599 | 0.5841 | 0.7431 | 0.9492 | 0.3084 | 0.7144 | ***0.0534*** | 0.4634 | ***0.1811*** | 0.2674 | 0.6882 | 0.9167 | 0.9717 |
| % starting weight - D7 | 0.4108 | ***0.1817*** | 0.4155 | ***0.0091*** | ***0.0587*** | ***0.1425*** | 0.9813 | ***0.0062*** | ***0.0209*** | ***0.0812*** | 0.6716 | 0.8164 | 0.5834 | 0.4359 |
| % starting weight - D8 | 0.7233 | 0.3180 | ***0.1179*** | ***0.0118*** | ***0.0486*** | ***0.1879*** | 0.5628 | ***0.0014*** | ***0.0136*** | ***0.1454*** | 0.9145 | 0.9509 | 0.4506 | 0.3135 |
| % starting weight - D9 | 0.3676 | 0.4294 | ***0.1076*** | ***0.0203*** | ***0.0992*** | ***0.0844*** | 0.4982 | ***0.0002*** | ***0.0032*** | ***0.1599*** | 0.8512 | 0.8397 | 0.4131 | 0.4464 |
| % starting weight - D10 | ***0.0090*** | 0.6912 | 0.2917 | ***0.0835*** | ***0.0286*** | 0.4658 | 0.6327 | ***0.0140*** | ***0.2001*** | 0.2786 | ***0.1680*** | 0.7355 | 0.7917 | 0.9000 |
| Day of lowest weight | 0.3815 | 0.8476 | ***0.1426*** | ***0.1130*** | 0.4531 | 0.2215 | 0.4623 | ***0.0013*** | ***0.0045*** | ***0.1506*** | 0.7092 | ***0.1419*** | 0.5138 | 0.5953 |
| Lowest weight reached | 0.2650 | 0.2469 | 0.2069 | 0.6501 | ***0.1149*** | 0.8742 | 0.6097 | 0.3364 | ***0.0165*** | 0.2501 | 0.2761 | 0.8172 | 0.8899 | 0.2907 |

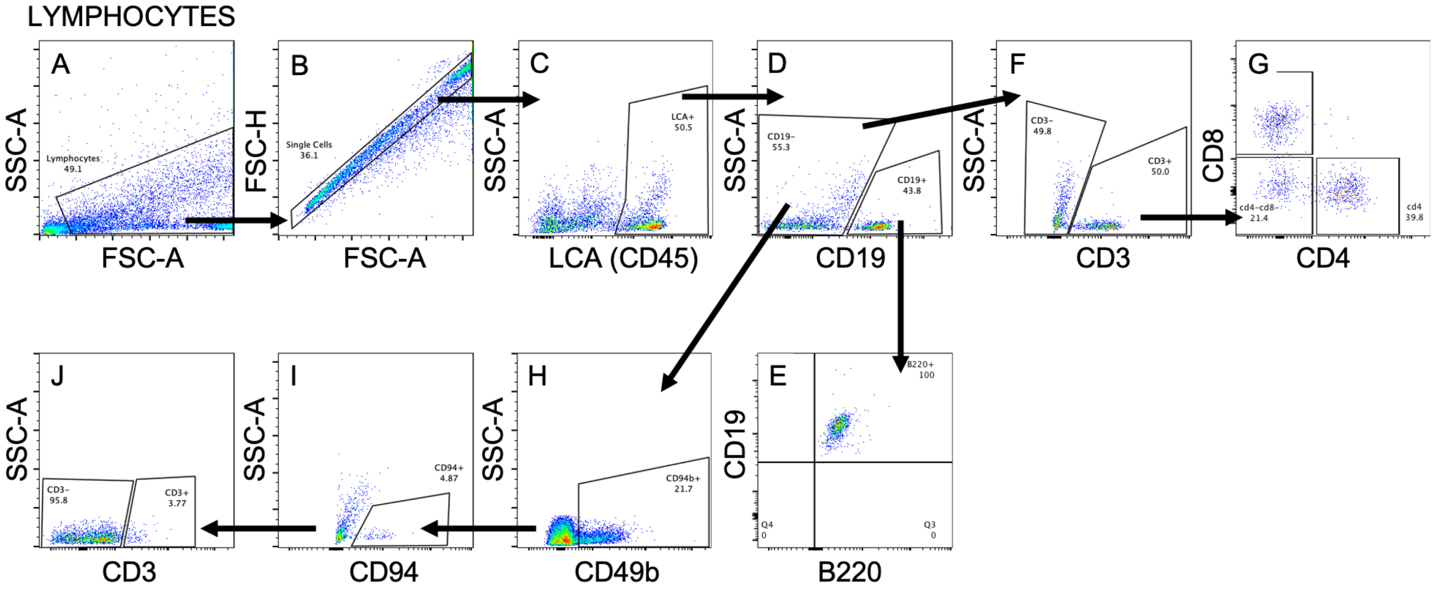


Supplemental Figure 1. Gating strategy to identify lymphocyte populations. All samples were first gated to eliminate debris, doublet, autofluorescent and non-hematopoetic cell populations. Next, events were gated to identify B cells (CD19+B220+; panel E), T cell subsets (CD3+CD4+, CD3+CD8+, or CD3+CD4-CD8-; panel G), and NK and NK T cells (CD49b+CD94+CD3- or CD49b+CD94+CD3+; panel J). NK and NK T cells were also gated through CD4 and CD8 negative populations to eliminate T cells from calculated frequencies.

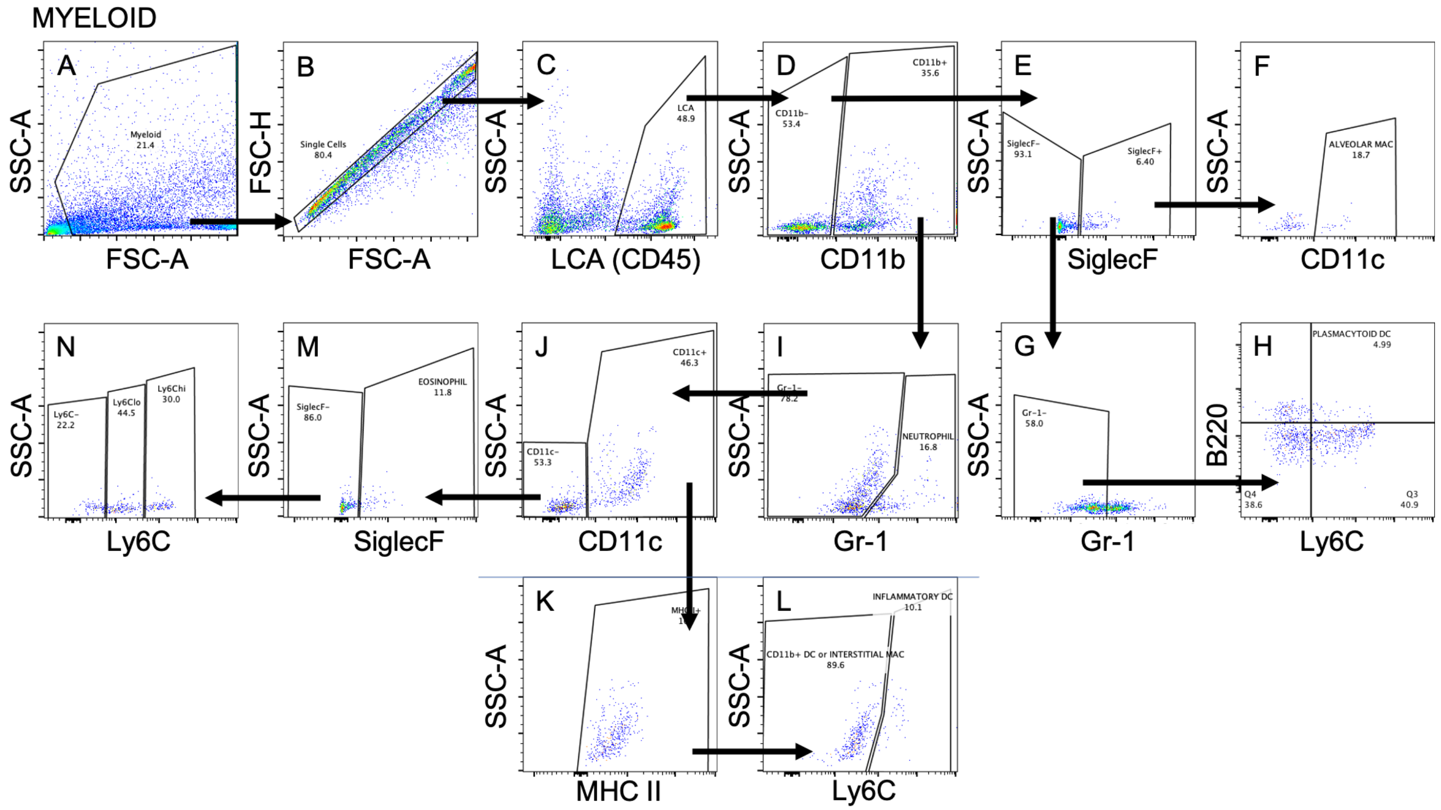


Supplemental Figure 2. Gating strategy to identify myeloid populations. All samples were first gated to eliminate debris, doublet and non-hematopoetic cell populations. Next, events were gated to identify alveolar macrophages (CD11b-SiglecF+CD11c+; panel F), plasmacytoid dendritic cells (CD11b-SiglecF-Gr-1-B220+Ly6C+; panel H), neutrophils (CD11b+Gr-1+; panel I), CD11b+ dendritic cells and/or interstitial macrophages (CD11b+Gr-1-CD11c+MHC II+Ly6C-; panel L), inflammatory dendritic cells (CD11b+Gr-1-CD11c+MHC II+Ly6C+; panel L), eosinophils (CD11b+Gr-1-CD11c-SiglecF+; panel M), Ly6C- monocyte/macrophage (CD11b+Gr-1-CD11c-SiglecF-Ly6C-; panel N), 'patrolling' monocyte/macrophage (CD11b+Gr-1-CD11c-SiglecF-Ly6Clo; panel N), and 'inflammatory' monocyte/macrophage (CD11b+Gr-1-CD11c-SiglecF-Ly6Chi; panel N).

DENDRITIC CELLS

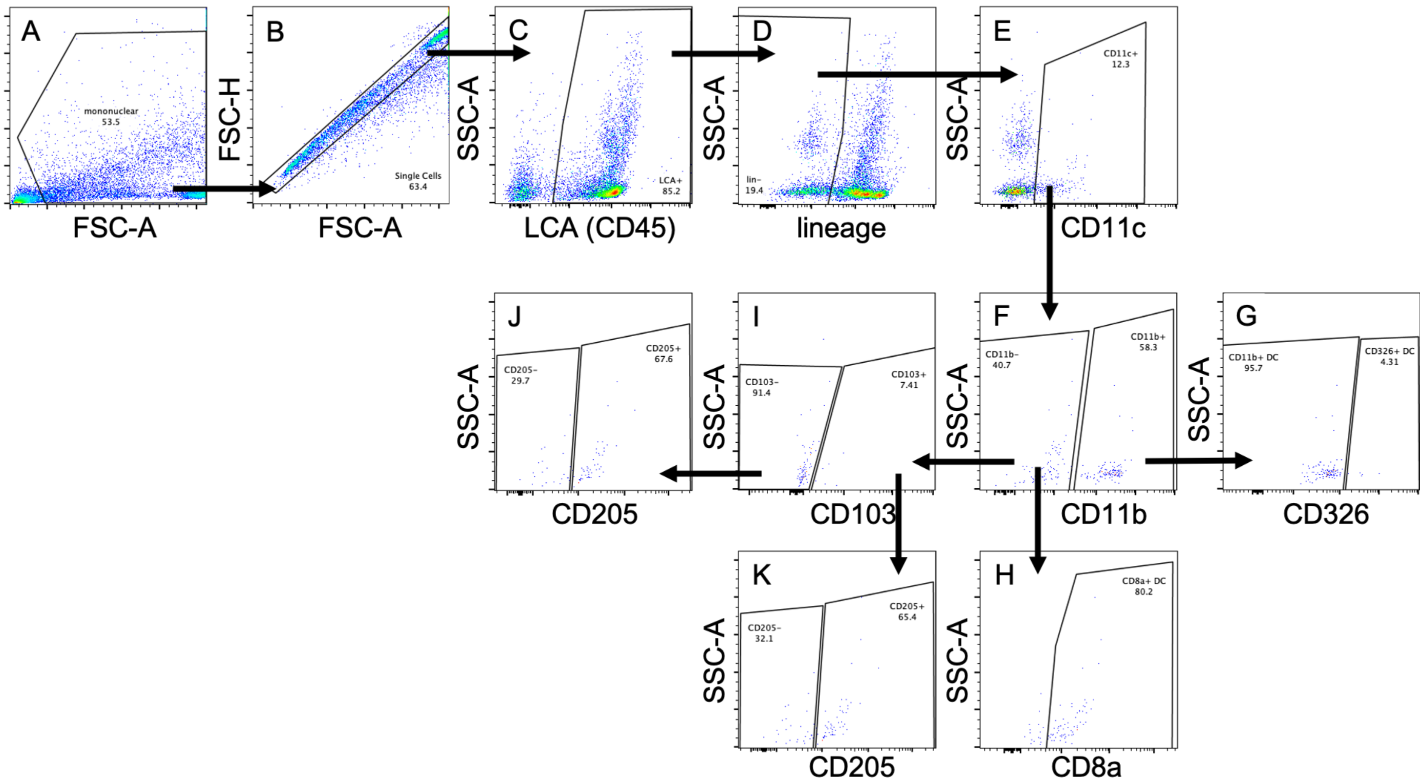


Supplemental Figure 3. Gating strategy to identify dendritic cell populations. All samples were first gated to eliminate debris, doublet, non-hematopoetic, lineage-, and CD11c- cell populations. Next, events were gated to identify CD11b+ dendritic cells (CD11c+CD11b+CD326-; panel G), CD326+ dendritic cells (CD11c+CD11b+CD326+; panel G), CD8a+ dendritic cells (CD11c+CD11b-CD8a+; panel H), CD103+CD205+ and CD103+CD205- dendritic cells (CD11c+CD11b-CD103+CD205+/-; panel K), and CD103-CD205+ and CD103-CD205- dendritic cells (CD11c+CD11b-CD103-CD205+/-; panel J).
